# Emergence of planar cell polarity from the interplay of local interactions and global gradients

**DOI:** 10.1101/2021.11.30.468750

**Authors:** Divyoj Singh, Sriram Ramaswamy, Mohit Kumar Jolly, Mohd. Suhail Rizvi

## Abstract

Planar cell polarity (PCP) – tissue-scale alignment of the direction of asymmetric localization of proteins at cell-cell interface – is essential for embryonic development and physiological functions. Abnormalities in PCP can lead to neural tube closure defects and misaligned hair follicles. Decoding the mechanism responsible for PCP establishment and maintenance remains a fundamental open question. While the roles of various molecules – broadly classified into “global” and “local” modules – have been well-studied, their necessity and sufficiency in explaining PCP and connecting their perturbations to experimentally observed patterns has not been examined. Here, we develop a minimal model that captures the proposed features of these modules – a global tissue-level gradient and local asymmetric distribution of protein complexes. Our model suggests that while polarity can emerge without a gradient, the gradient can provide the direction of polarity and maintain PCP robustly in presence of stochastic perturbations. We also recapitulated swirling patterns seen experimentally and features of domineering non-autonomy, using only three free model parameters - protein binding rate, concentration of proteins forming heterodimer across cell boundaries and gradient steepness. We explain how self-stabilizing asymmetric localizations in presence of tissue-level gradient can lead to robust PCP patterns and reveal minimal design principles for a polarized system.

## Introduction

Epithelial tissues form the outermost layer of most organs including skin. They not only function as a protective layer for the inner tissues but also take part in several physiological functions including absorption, secretion, excretion and filtration [1]. Epithelial cells usually manifest two major types of polarity: *Apico-basal polarity* is asymmetry in the distribution of cellular components and functions between the two surfaces of an epithelial sheet [2], and *planar cell polarity* (PCP) refers to organization in a direction parallel to the plane of the epithelial sheet [3]. PCP is an evolutionarily conserved phenomenon which is required for early developmental events such as neural tube closure and later in the formation of functional organs [4]. It has been a topic of extensive scientific investigation, especially in the model organism *Drosophila melanogaster*. In *Drosophila*, PCP has been investigated to understand its molecular mechanisms and the fallouts of its disruptions in the physiology of several organs, such as hair bristles on wing, abdomen and ommatidia in eye [5].

Disruption in PCP can lead to a variety of developmental abnormalities such as neural tube defects, idiopathic scoliosis, skeletal dysplasia manifested as short limbs and craniofacial anomalies [3] and tracheal ciliary malfunction [6]. PCP is essential also for auditory and vestibular function of the vertebrate inner ear. It determines how mechanosensory hair cells sense sound or motion [7]. Thus, defects in PCP have been associated with hearing loss [8]. Therefore understanding the multi-scale dynamics of PCP establishment and maintenance is a question of fundamental importance.

The molecular players involved in the establishment of PCP in various organisms have been well studied. Though the interactions among the proteins involved in PCP are quite intricate and not very precisely demarcated, they have been broadly categorized into two groups, also known as *modules*: Global and Core [3].

The global module contains two atypical cadherins, Fat (Ft) and Dachsous (Ds), the kinase Four-jointed (Fj), and the myosin Dachs(D) [9]. Ft and Ds from adjacent cells can form heterodimers at cell-cell interface. Their affinity for each other can be modulated by Fj [10–12]. Dachs co-localises with Ds and generates active forces at the apical end, thus regulating organ shape [13]. Proteins of global modules are usually present in tissue-level expression gradients [14–16]. These tissue-level gradients are known to provide the global cue to PCP machinery which helps alignment of the PCP with the axis of the organ [5].

On the other hand, the core or local module consists of Frizzled (Fz), Flamingo (Fmi), Van Gogh (Vang), Prickle-spiny-legs (Pk), Dishevelled (Dsh) and Diego (Dgo) [17] which interact with one another, some intercellularly and some intracellularly, by forming complexes. Similar to Ft-Ds heterodimers across a cell-cell boundary, Fmi from two neighbouring cells form complexes. The trans-membrane protein Fz recruits the cytosolic proteins Dsh and Dgo, and localises in proximity of Fmi in one of the two cells. Another trans-membrane protein Vang recruits cytosolic protein Pk close to Fmi on the other cell. These interactions within the core module have been argued to be sufficient for establishing a local order (polarity in two adjacent cells) [3, 18].

An understanding of PCP crucially requires establishing the relative contributions of global agencies such as gradients and local interactions like heterodimer formation in the emergence and maintenance of polarity [22, 23]. Most models have thought of global interactions as a means for only providing directional cues [24]. Local interactions such as the formation of heterodimers have been considered as an output of the formation of polarity and used as a readout of polarization [25]. In principle, global fields such as tissue-level gradient can of course help *maintain* polarity in the presence of perturbations but, if inter-cellular interactions are weak, they can *create* polarity on their own only to an extent proportional to their own magnitude. Strong enough local interactions can cooperatively give rise to a strong susceptibility for macroscopic polarity. A global field can then produce a disproportionate tissue-scale polarization. How redundant or synergistic the roles of these two features – asymmetric localisation and tissue-level gradient – are remains to be decoded.

Given the complex interactions between molecules within and across modules, mathematical modelling has been an essential tool to account for experimental abnormalities and make further testable experimental predictions. Despite the two-dimensional nature of epithelial tissue, most mathematical models of the PCP have been formulated for one dimension (for example [26–29]), with exceptions where the biological question studied required a two-dimensional framework [25, 30–32]. There are broadly two classes of models for PCP. One class takes into account the mechanistic details of the system and explains examples of PCP establishment and its disruption in specific organs (see [33] for review). While they provide useful inputs for the experimental contexts for which they are formulated, their generalizability across parameter sets or biological contexts is limited. The other class are phenomenological models that apply to a wider variety of cases and parameter ranges because they ignore some mechanistic details but account for the pattern formation [31].

Here, we integrate the strengths of both these kinds of models in the context of the global module of PCP. We incorporate minimal mechanistic relations with the goal of capturing general dynamical properties of the system, together with focus on specific interesting experimental observations. Towards this, we focus on the inter-cellular interactions of Ft and Ds to form hetero-dimer complexes and look at the effect of hetero-dimer formation on asymmetric localization of these proteins on cell membranes. We have developed a minimal model of the asymmetric enrichment of Ft and Ds in one and two dimensional frameworks with only three free parameters. At the microscopic level we consider the interaction between two proteins at the cell-cell interface, in contrast to other works in which a presumed inter-cellular interaction between inter-cellular hetero-dimers is considered [27, 30].

With this minimal model we recapitulate some of the known PCP results such as the establishment of PCP without any global cue [34]. We further demonstrate the dual role of the tissue-level expression gradients in PCP establishment and its coordination with tissue axis. We also test the robustness of the PCP in the minimal model in the presence of static and dynamic noises representing cell-to-cell variations in expression levels of the proteins and thermal fluctuations in protein binding kinetics, respectively. Finally, we also explain non-homogeneous PCP patterns on deleting specific proteins from a region and make predictions which can be tested experimentally.

## Materials and Methods

### Basic assumptions

We assume epithelial tissue to be a non-motile confluent layer of cells. For simplicity, we ignore the three dimensional organ-specific epithelial architecture and assume identical apical-basal geometry of the cells and a flat planar tissue, and hence a purely two-dimensional description, which is adequate for describing planar cell polarity. This is a fair simplification and has remained a common feature across all the PCP modeling approaches [25, 30, 31]. In order to keep the analysis limited to PCP we also do not take into account any cellular rearrangements, cell movements or remodeling in our model. This simplification is based on the assumption that we are focusing on the late stages of PCP establishment where cellular rearrangements have already taken place. We assume that the protein translation and degradation happens at very slow rate (large time scale) and protein diffusion inside the cell takes place at a very fast rate (small time scale). We assume the PCP dynamics to take place at a rate somewhere between the two. Therefore, we do not consider any change in the total protein concentration in a cell in this work.

### Protein binding and transport kinetics

In order to build the model we focus on the two members of the global module of PCP, Fat (or Ft) and Dachsous (or Ds), atypical cadherins which interact across the membranes of two adjacent cells (inter-cellular interaction). Both Ft and Ds, if present in the cytoplasm, can bind to the cell membrane and, if bound to the cell membrane, can detach and enter the cytoplasm. We assume within our model that in an ‘isolated’ cell (a hypothetical situation where inter-cellular interactions are ignored) the attachment and detachment of these cadherins follow the law of mass action and the kinetic rates of attachment and detachment are *α* and *β*_0_, respectively. We make the further simplifying assumption that these rate constants are the same for both proteins but the argument can easily be generalized to disparate rate coefficients.

Finally, we take inter-cellular protein interaction at the cell membrane into account in the following manner. These proteins are cadherins, and they have been shown to form Ft-Ds heterodimers across cell membranes. Therefore, when an Ft (or Ds) is bound to the cell membrane its rate of detachment goes down due to its interaction with the Ds (or Ft) from neighboring cell. A rate of detachment of the form

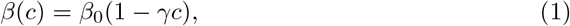

where *c* is the concentration of Ds or Ft, as the case may be, at the membrane of the neighboring cell, should be adequate for small *c. γ* here is the facilitation parameter for repressing the unbinding of Ft from the cell membrane by Ds of neighboring cell (or vice-versa) and forming a heterodimer.

With this simple biochemical kinetic assumption we now move on to the formulation of the dynamics of PCP in the tissue. First, we will describe it for a one dimensional tissue.

### One dimensional model

In the 1-D model, we assume a single row of cells. The total concentrations of Ft and Ds in cell are held constant thanks to the assumption mentioned previously.

Since in 1D each cell has only two ends, the concentrations of membrane bound Ft at either sides of the cells are denoted by *f*_*l*_(*i*) (amount of *Ft* on left edge of the cell at location *i*), *f*_*r*_(*i*) (amount of *Ft* on right edge of the cell at location *i*). Similarly, for Ds these quantities are denoted by *d*_*l*_(*i*) and *d*_*r*_(*i*).

The equations describing the concentrations of membrane bound proteins are

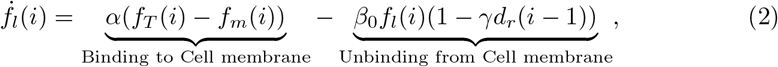

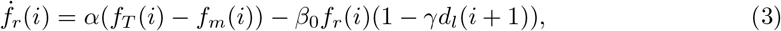

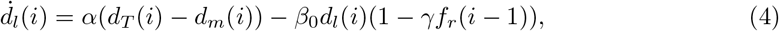

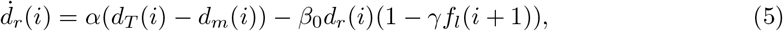

where *f*_*m*_(*i*) = *f*_*l*_(*i*) + *f*_*r*_(*i*) and *d*_*m*_(*i*) = *d*_*l*_(*i*) + *d*_*r*_(*i*) are the total membrane-bound protein in the *i*^th^ cell. The two terms on the right-hand sides of the equations represent the rates of protein binding and unbinding (which is dependent on the protein concentrations of other protein in the neighboring cell), respectively. Here, *f*_*T*_ (*i*) and *d*_*T*_ (*i*) stand for the amount of total (cytoplasmic+membrane bound) Ft and Ds present in the cell and can be dependent on the cell position in the tissue.

In this work, we define the cell polarity in terms of the degree of asymmetric localization of the two proteins in the cell. We write the asymmetry or polarity of the two proteins in the *i*^th^ cell as the difference between the protein concentrations on right and left edges of the cell, that is

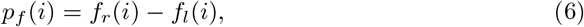

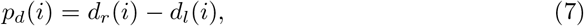

Finally, using these definitions of protein polarity we define the overall polarity

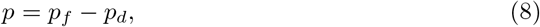

the observable of central interest in this work.

To simulate the 1D model, we considered an array of 500 cells and numerically solved the system of ordinary differential equations using the function solve ivp from the scipy package(version 1.6.2) in Python with adaptive time-stepping.

### Effect of noise in the system

#### Noise in protein kinetics

To study the effect of noise in the system due to protein kinetics, we write stochastic equations for the system. Note: We do not expect our *one-dimensional* models with noise included to yield PCP in the limit of infinite system size, but they will give rise to coherent polarization on scales that grow as the noise intensity is decreased. For the stochastic simulations for the 1D case, we modify the equations by adding zero-mean Gaussian white noise terms of small magnitudes, leading to the Langevin equations

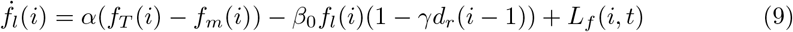

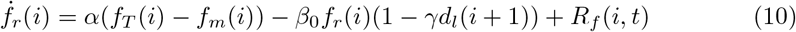

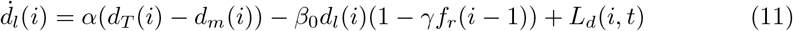

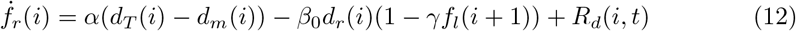

where the noise terms (such as *L*_*F*_ (*i, t*)) are uncorrelated in time *t* and position *i* of the cell in the tissue, and with respect to location *L* and *R* of the cell edges, and protein type *Ft* and *Ds*, that is

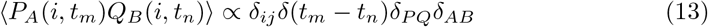

where *P* and *Q* can take values *L* and *R, A* and *B* label the protein type (Ft or Ds), and the *δ* symbols are Dirac or Kronecker deltas depending on whether their argument is a continuous or discrete variable. Here, we are looking at small fluctuations in the levels of membrane-bound proteins. In order to implement this we select a random number for each cell edge *T* (*i, P, A, t*_*m*_) ∈ 𝒩 (0, 1) from a standard Normal distribution with mean *μ* = 0 and standard deviation 1. Using this, we write the noise term in the equation as

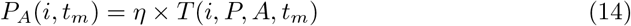

where *η* represents the amplitude of the noise or the magnitude of stochasticity in the system. We present results for the polarization averaged over 50 realizations of the noise.

#### Static spatial noise in total protein concentration of Ft and Ds

Further, even for a uniform tissue level protein expression no two cells are expected to have identical protein levels. Therefore, we have also considered small cell-to-cell variations in the total protein levels:

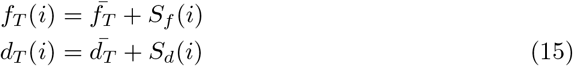

where 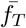 and 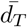 are the average protein levels in the cells of the tissue and the noise terms *S*_*f*_ (*i*), *S*_*d*_(*i*) are uncorrelated along position *i* of the cell in the tissue, that is

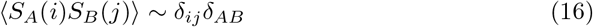

where the labels *A, B* can be *f* or *d*, and *i* and *j* stand for the position of the cell. To implement noise in total protein concentration, we selected a random number *T* (*A, i*) ∈ 𝒩 (0, 1) from a Standard Normal distribution with mean 0 and standard deviation 1. Using this, we write the noise term in the equation as

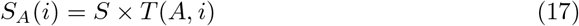

where *S* represents the amplitude of the noise or the magnitude of stochasticity in the system. Our results for the polarization are averaged over 50 realizations of the noise.

### Two-dimensional model

For the 2D model, we used a lattice of immotile cells formed of regular hexagons with each cell having six neighbors as opposed to two in 1D. Following same approach as in 1D, we write equations for dynamics of protein concentration for a cell at **x**:

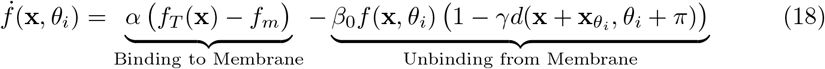

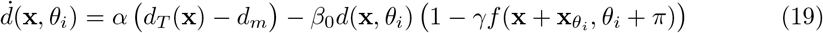

where

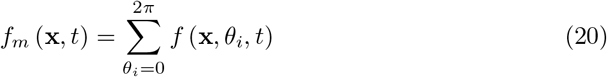

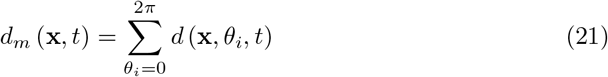

are the total membrane-bound protein concentrations in the cell. Here we sum over the protein concentration at the 6 angles *θ*_*i*_ = 0 = 2*π, π/*6, *π/*3, *π/*2, 2*π/*3 and 5*π/*6. (6 parts of discretised cell membrane). *f*_*T*_ (**x**, *t*) and *d*_*T*_ (**x**, *t*) are the total (cytoplasmic+membrane bound) protein levels in the cell located at **x**, and 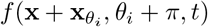 represents the protein *Ft* in the neighboring cell.

Similar to the one-dimensional model, we can define the planar cell polarity (PCP) of a protein in a cell in terms of the asymmetric distribution of that protein on the cell membrane, leading to the vectors

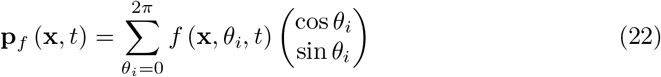

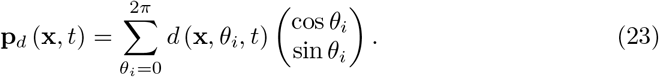

characterizing the local orientational asymmetry in the distributions of Ft and Ds. Finally we define

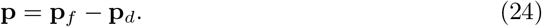

To simulate the 2D system, we take a 2D hexagonal lattice of 50 × 50 cells and solve the equations for the Ft and Ds dynamics at cell boundaries using Euler’s method.

### Parameter values

To understand the system analytically and reduce the modelling parameters, we non-dimensionalize the system using the following characteristic quantities

1. *τ* = 1*/β*_0_ for time,
2. 1*/γ* for protein levels.

This results in just three free parameters in the system-Protein binding rate (*α*), total protein concentrations *f*_*T*_ and *d*_*T*_. These parameters along with the initial conditions can control the wide variety of patterns seen in PCP. Ft and Ds, being membrane proteins, do not stay in the cytoplasm and bind to the membrane at a fast rate which results in large *α*. We have kept the value of non-dimensionalized *α* fixed at 10 in the results shown unless specified otherwise. The total number of time (non-dimensionalized) for 1D simulations are 2 × 10^4^ and for 2D simulations are 5 × 10^4^. The protein concentrations are used as a control parameter to study the nature of polarization and the effects of protein gradient expressions, protein loss and gain.

## Results

### Polarity can emerge even in the absence of gradient in protein expression

In PCP, tissue-level gradients of Ft and Ds are well documented [21, 35–37], but their exact role and the mechanism of their action is not completely understood. Thus, we first ask whether polarity can emerge in absence of any tissue-level gradient and noise in 1D model. Here for simplicity we have kept the levels of Ft and Ds in each cell equal to *ρ*, referred to as total protein concentrations (membrane+cytoplasm), that is *f*_*T*_ = *d*_*T*_ = *ρ*. The results remain valid even if these levels are unequal. When *f*_*T*_ ≠ *d*_*T*_, the polarization magnitude of Ft and Ds become unequal.

We observe that with an increase in *ρ* the total concentrations of membrane bound Ft and Ds increase equally, that is *f*_*l*_ + *f*_*r*_ = *d*_*l*_ + *d*_*r*_ := *ρ*_*m*_. However, there is a threshold of total protein concentration, *ρ*_*c*_, after which an increase in *ρ* results in abrupt but continuous (Fig. S2) increase in the membrane bound protein levels (**Fig** 2C). Below the threshold protein concentration, both the proteins localise on both sides of the cell in equal amount. Therefore there is no polarization of the cells (**Fig** 2D and **Fig** S1). But once we cross the threshold concentration, that is *ρ > ρ*_*c*_, Ft gets bound to one side of the cell and Ds on the other side, resulting in non-zero polarization (**Fig** 2 D and **Fig** S1). The direction of polarization here is determined by the initial conditions. The presence of threshold is due to the inter-cellular interactions and heterodimer formation between the proteins across the membranes of two neighboring cells(**Fig** 2 E). We note that the transition from unpolarised state to polarised state is continuous one S2.

**Figure 1.**
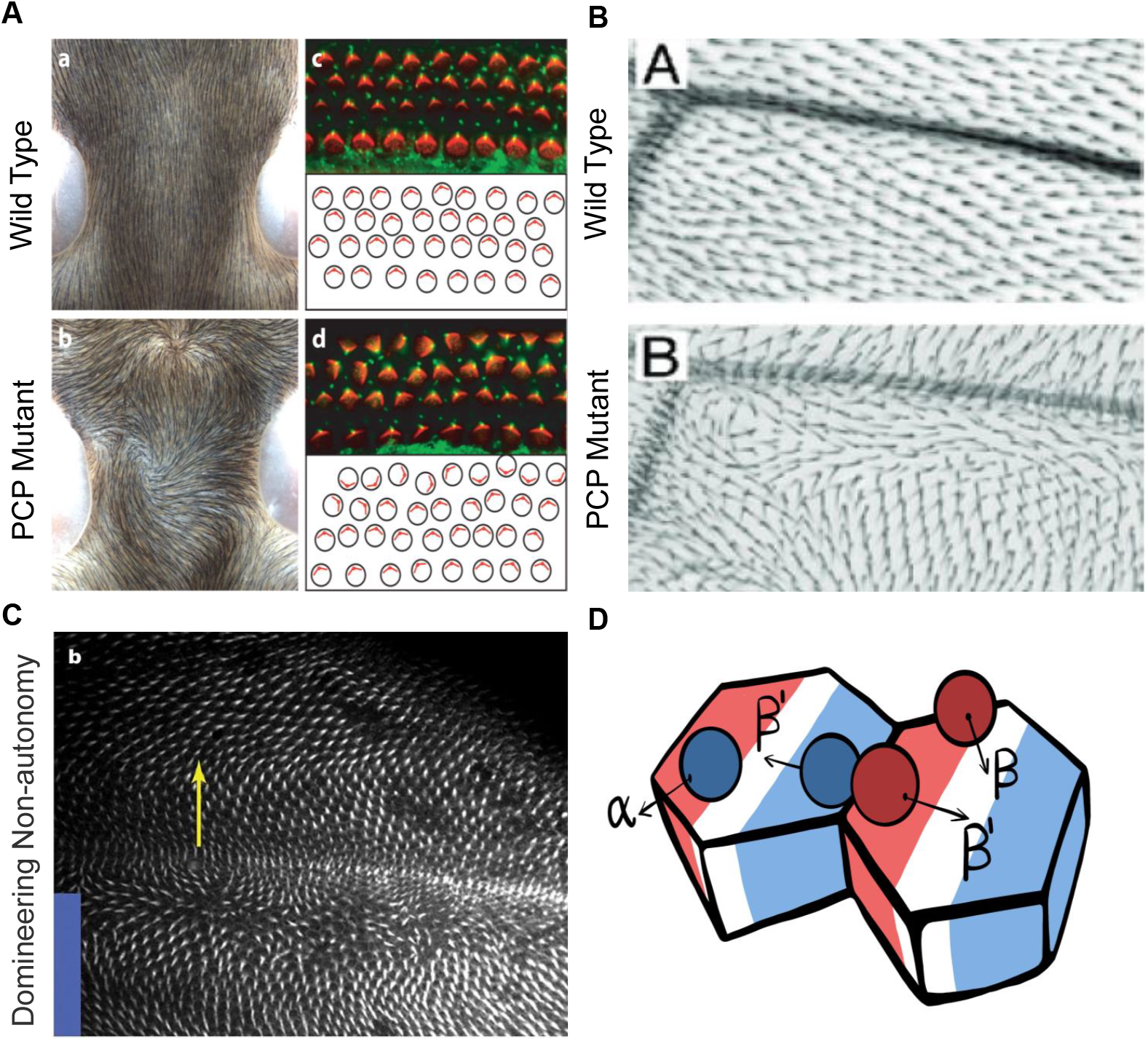
Introduction to Planar cell polarity and the model. **A)** Examples of PCP in mammals (mouse skin and inner ear) taken from [19]. **B** Example of PCP in wing of *Drosophila melanogaster* taken from [20] **C)** Experimental image showing domineering non-autonomy taken from [21]. **D)** Schematic to depict the protein binding and unbinding kinetics and heterodimer formation.

**Figure 2.**
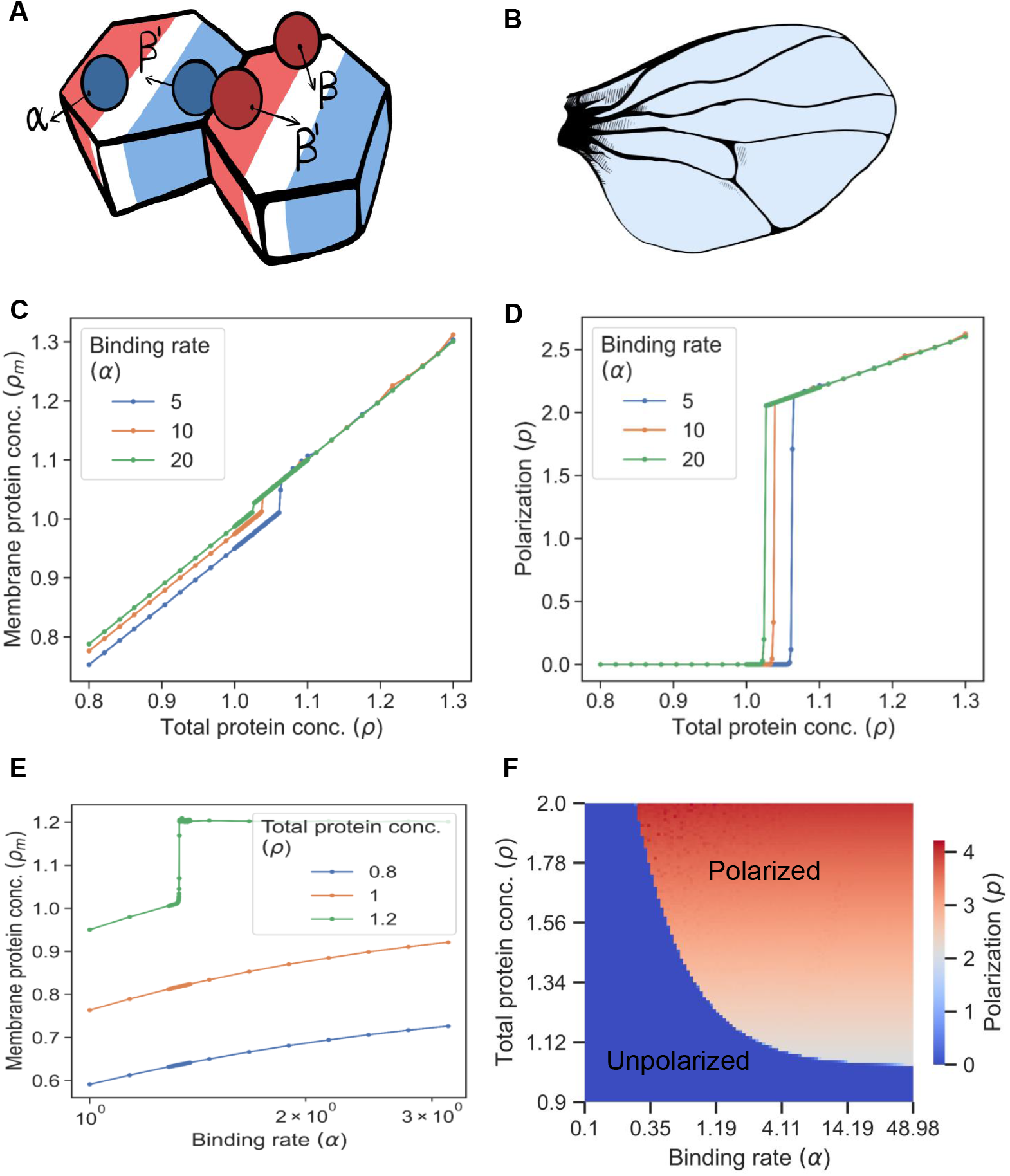
Emergence of polarity due to inter-cellular interactions in absence of any tissue level gradient in expression. **A)** Schematic describing the binding and unbinding of the proteins at the membrane and its regulation by heterodimer formation. *α* is the binding rate of the protein towards the cell membrane and *β*_0_ is the unbinding rate. *β* is the reduced unbinding rate if the heterodimer is formed. **B)** Schematic to show uniform expression levels of both proteins across the tissue. Note that the results in this figure are from the one-dimensional model. **C)** Membrane protein concentration (of Ft and Ds) (*ρ*_*m*_) at the steady state against total protein concentration (*ρ*). **D)** Plot of polarization magnitude (*p*) against total protein concentration (*ρ*). Tissue gets polarized above a threshold of total membrane concentration. **E)** Plot of membrane protein concentration (of Ft and Ds) (*ρ*_*m*_) at steady state as a function of protein binding rate (*α*) at different total protein concentrations (*ρ*). **F)** Phase diagram of polarization (*p*) as a function of binding rate (*α*) and total protein concentrations (*ρ*) shows the existence of a polarized (red) and unpolarised state (blue).

Next we studied the dependence of the membrane bound protein concentrations (*ρ*_*m*_) as a function of the protein binding rate (*α*). Since the threshold protein concentration can depend on the binding rate *α* we generated a phase diagram of polarization as a function of protein binding rate, *α* and total protein concentration (*ρ*). The phase diagram shows the existence of a polarized state (*p* ≠ 0) and an unpolarised state (*p* = 0). The boundary separating the two states shows that with an increasing *α* the threshold concentration of proteins required for the polarization decreases.

In order to get an intuitive understanding of the establishment of polarization above the critical protein concentrations, we looked at the continuum limit of our discrete model of 1D epithelium (see Supporting Information for details) [38]. In the continuum limit the size of the single cell is taken to be much smaller as compared to the length scale of the tissue. In this limit we get a few partial differential equations in place of the large number of ordinary differential equations (2)-(5) we have described earlier. In the case of uniform protein expression levels, we get the following equation for the polarization (in non-dimensional form)

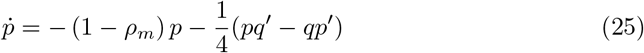

where *p* is the cell polarity as described in equation (8) and *ρ*_*m*_ is the total concentration of the membrane bound proteins. Eq. (25) tells us that if *ρ*_*m*_ *>* 1 the state of zero cell polarity is unstable, and a nonzero value of polarization will set in. The equation says that the cell polarity (*p*) will increase indefinitely, but this is simply a limitation of the linearised form (25). As soon as at *ρ*_*m*_ = *p/*2 (which is also seen in Fig. 2E-F) the two proteins get localized on two edges of the cell and further change in concentrations stops. Thus, we show that the inter-cellular interactions, in the form of heterodimer formation between Ft and Ds, are sufficient for the emergence of polarity in the absence of any tissue level expression gradient. Further nontrivial properties of the polarized state, including a possible destabilizing response to spatially varying perturbations, are discussed in our study of the continuum model [38].

### Stochasticity results in partial loss of tissue level coordination in PCP

We have seen how spontaneous polarization of tissue can emerge in a completely deterministic treatment in which we model the dynamics of average occupancies. Binding and unbinding are, however, intrinsically stochastic and, more generally, noise is ubiquitous in biological systems [39].

In the context of PCP, the noise can arise from a variety of sources with a range of timescales. One source of noise are cellular processes like protein production and degradation, and cell division, growth, birth and death which can affect the total protein concentration of Ft and Ds in each cell. These can lead to variation in protein concentrations across the tissue. We have examined the impact of such stochastic effects on the polarization in the tissue.

We have incorporated static or “quenched” noise in the equations as described earlier in (15). For simplicity, we have assumed the noise amplitude to be small compared to the average concentration *ρ* of protein in each cell, in which case it can be taken as purely additive. We start with calculating the average polarization in the tissue (*p*) (averaged over multiple simulations) in the system for a range of noise amplitude (*S*) (**Fig** 3 A). We observe that the tissue starts losing polarization if the noise level exceeds a threshold.

**Figure 3.**
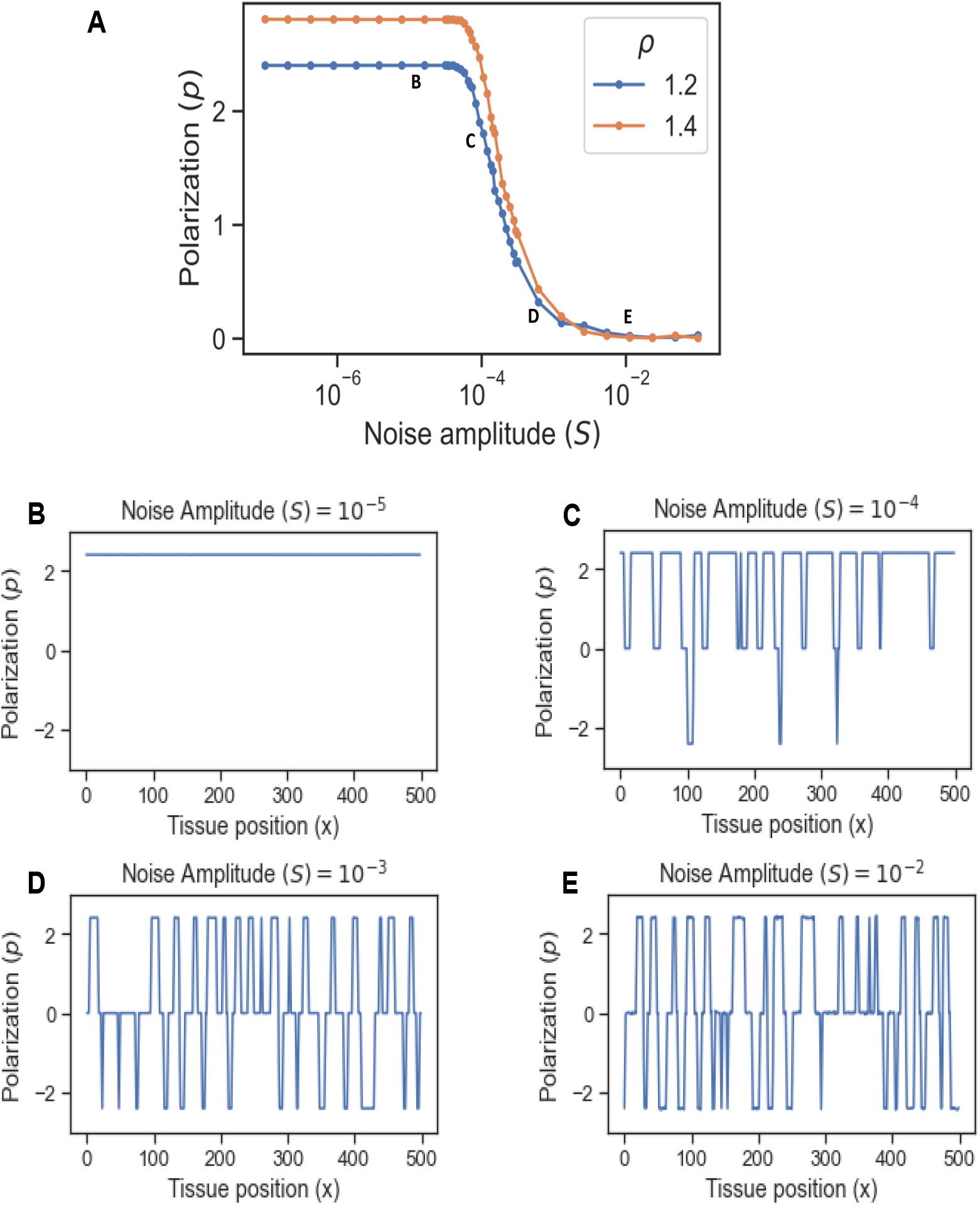
Effect of noise in total protein concentrations on polarization. **A)** Polarization (*p*) as function of noise amplitude (*S*) for different total protein concentrations (*ρ*). **B)** Polarization (*p*) in the tissue when the level of noise is negligible (*S* = 10^*−*5^). **C)** Polarization at high amplitude of noise (*S* = 10^*−*4^), some parts of the tissue are aligned in the opposite direction **D)** At higher noise amplitude (*S* = 10^*−*3^), the average polarity of the tissue reduces **E)** At very high noise amplitude (*S* = 10^*−*2^), the tissue loses overall polarization as different parts of the tissue are aligned in opposite direction.

The other source of noise in PCP arises from the nature of the protein binding and unbinding from the membrane or the inter-cellular interactions at the cell-cell interface. We incorporated this dynamical noise in the equations as described earlier in Methods ((10)-(12)). Again, we have considered the noise amplitude to be not very large as compared to a scale determined by *β*_0_*/γ*, the characteristic rate of the protein kinetics in this system. We again calculated average polarization in the tissue (*p*) (averaged over 50 simulations) in the system for a range of noise amplitude (*η*) (**Fig** S3 A). We observe behaviour similar to that seen for the effect of the noise in total protein concentration. Once the noise amplitude (*η*) is higher than a threshold level, the tissue starts losing polarization.

We ask next whether noise results in a reduction in the magnitude of the polarization at the scale of a cell or, instead, leaves the magnitude locally unchanged but randomizes its direction on larger scales so that the spatial average is zero.

To address this, we looked at the polarity of each cell in the tissue for different amplitudes of static (*S*) and dynamic (*η*) noise. Note that these results are for one instance of simulation.

When the noise amplitude is small (*S* = 10^*−*5^), the tissue remains polarized uniformly. The direction of polarity of the tissue is determined by the initial conditions (**Fig** 3 B). As we increase the noise amplitude (*S* = 10^*−*4^), some of the cells are polarised in opposite direction (**Fig** 3 C). On further increasing the amplitude of noise (*S* = 10^*−*3^), the overall polarity reduces due to patches of tissue polarising in opposite directions (**Fig** 3 D). The tissue as a whole eventually loses polarity if the noise amplitude is too high (**Fig** 3 E) because of cells polarizing in opposite directions randomly. For the range of noise strengths explored, the individual cells remain polarized. Similar results are seen when noise is added to the protein kinetics equations (**Fig** S3 B-E) These observations show that over the parameter range studied the noise predominantly influences the overall coordination of the the polarity in the tissue with minimal effect on PCP of individual cells.

We close this subsection with a word of caution. As we remarked above, any nonzero noise should destroy order (i.e., spontaneous macroscopic polarity) in the 1D models. So in the 1D models what might appear to be an ordered phase is simply a low-noise state in which the correlation length, while finite, exceeds the system size.

### Swirling PCP patterns are possible in the absence of gradient in expression

Several experimental works [21] report swirling patterns in the PCP, especially when the global cue, that is, the tissue level expression gradient, is absent [13]. We therefore explore the occurrence of non-uniform polarization of the tissue within our model. Swirls being a two-dimensional phenomenon, we study the case of a 2D hexagonal lattice, which is a fair approximation to the actual epithelial geometry. We start with the case where the total protein concentration of both the proteins is uniform throughout the tissue and is equal to *ρ* (**Fig** 4 A). Similar to the 1D case, we observe that the tissue gets polarized if the total protein concentration is above a threshold value (**Fig** 4 B). See [38] for a more detailed analysis.

**Figure 4.**
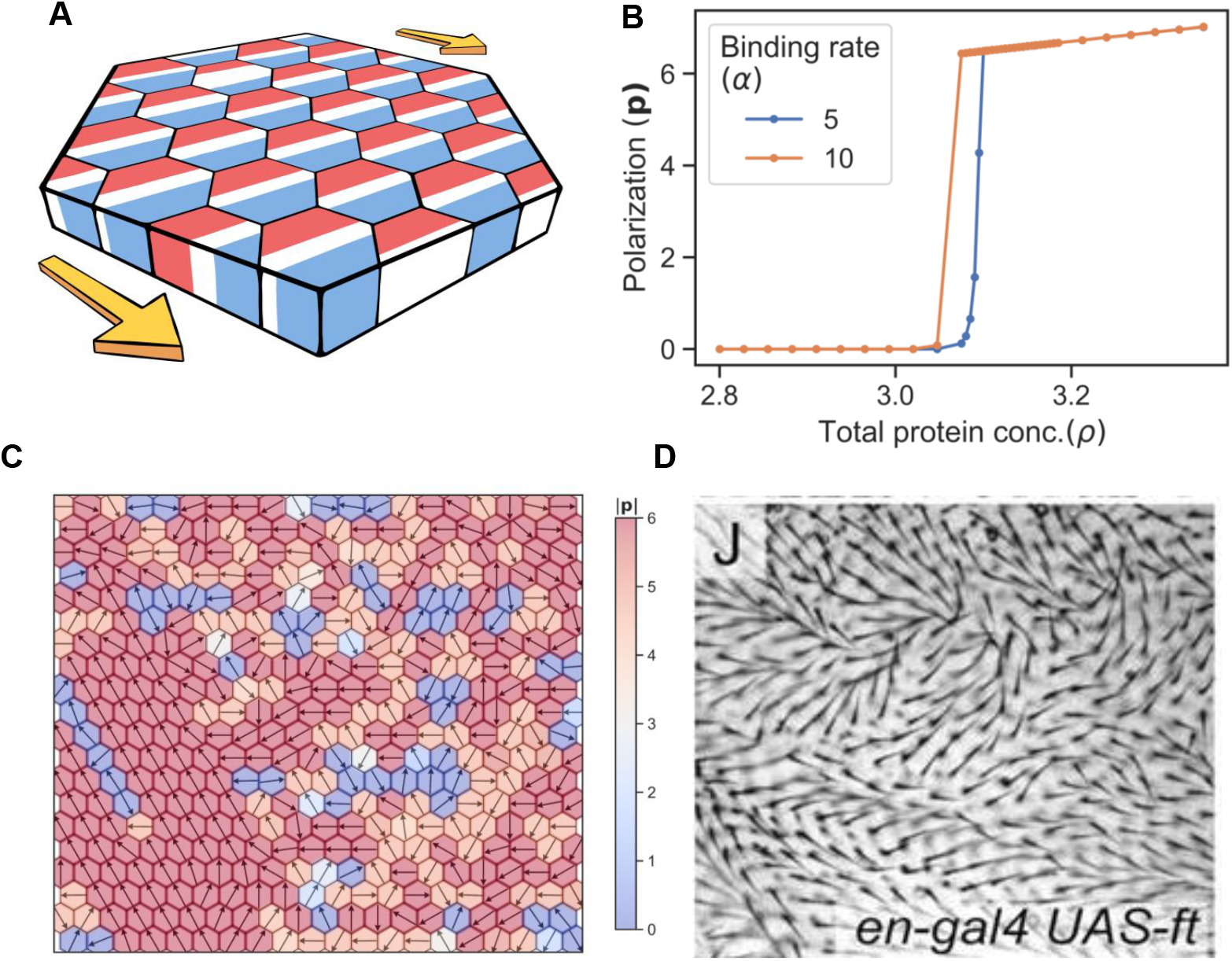
Polarization in 2D lattice for uniform protein expression of Ft and Ds: **A)** Schematic to describe the model of uniform protein expression of Ft and Ds in the 2D layer of tissue. **B)** Polarization (|**p**|) as a function of total protein concentration (*ρ*). The system starts getting polarized above a critical value of total protein concentration. **C)** Swirling patterns are observed when noise is introduced in the total protein concentration (*S* = 10^*−*8^) **D)** Experimental image of global mutant taken from Axelrod et al. 2020 [40] which shows and swirling patterns **E)** Patches of tissue polarized in different direction when the amplitude of noise in total protein concentration is higher(*S* = 10^*−*3^). **F)** Experimental image taken from [42] depicting a case where we observe patches of tissue polarized in different directions.

Noise (both due to variation in total protein concentration and protein kinetics) in our two dimensional model leads to non-homogeneous polarization in the tissue. We observe a variety of patterns some of which have been famously observed in PCP [13]. First we observe aster and swirling patterns where polarization (**P**) in multiple cells either originate or terminate, or rotate around a point, respectively (**Fig** 4 C,**S Video** 1, **S Video** 2). These patterns are static in time once the cells are polarized. The amplitude of noise in total protein concentration is very small for this case (*S* = 10^*−*5^). This is an interesting finding and accounts qualitatively for the swirling patterns observed in experimental systems by Axelrod et al. (2020) [40] as shown in (**Fig** 6 D) or at least suggests that they are a natural outcome of PCP physics. These types of swirling patterns have been studied recently in out of equilibrium disordered active systems [41].

Next, we observe a case where polarization is in different directions in patches (**Fig** 6 E). The amplitude of noise in total protein concentration is higher as compared to earlier (*S* = 10^*−*1^). This captures the experimental findings observed by Matakatsu et al. (2004) [42]. This shows that in 2D the PCP direction need not be along a preset direction, such as the tissue axis.

### Gradients in protein concentrations facilitate PCP and provide global cue

As mentioned earlier, the proteins of global module of PCP are expressed as a gradient at the tissue level [13]. These gradients have been studied extensively in the context of the wing development in *Drosophila* (**Fig** 5A). Therefore, we also investigated the effect of the expression gradients of the proteins on PCP establishment and its alignment with tissue axis. For simplicity, we start with studying the effects of a linear gradient of Ft. Gradient of Ds have similar effects. The choice of a linear gradient is inspired by experimental studies [21, 42, 43] that demonstrated the presence of gradients that are either linear [12] or can be represented by piece-wise linear approximations. Several other modelling works [26, 27] have also taken gradients to be linear in the position of cell in the tissue.

**Figure 5.**
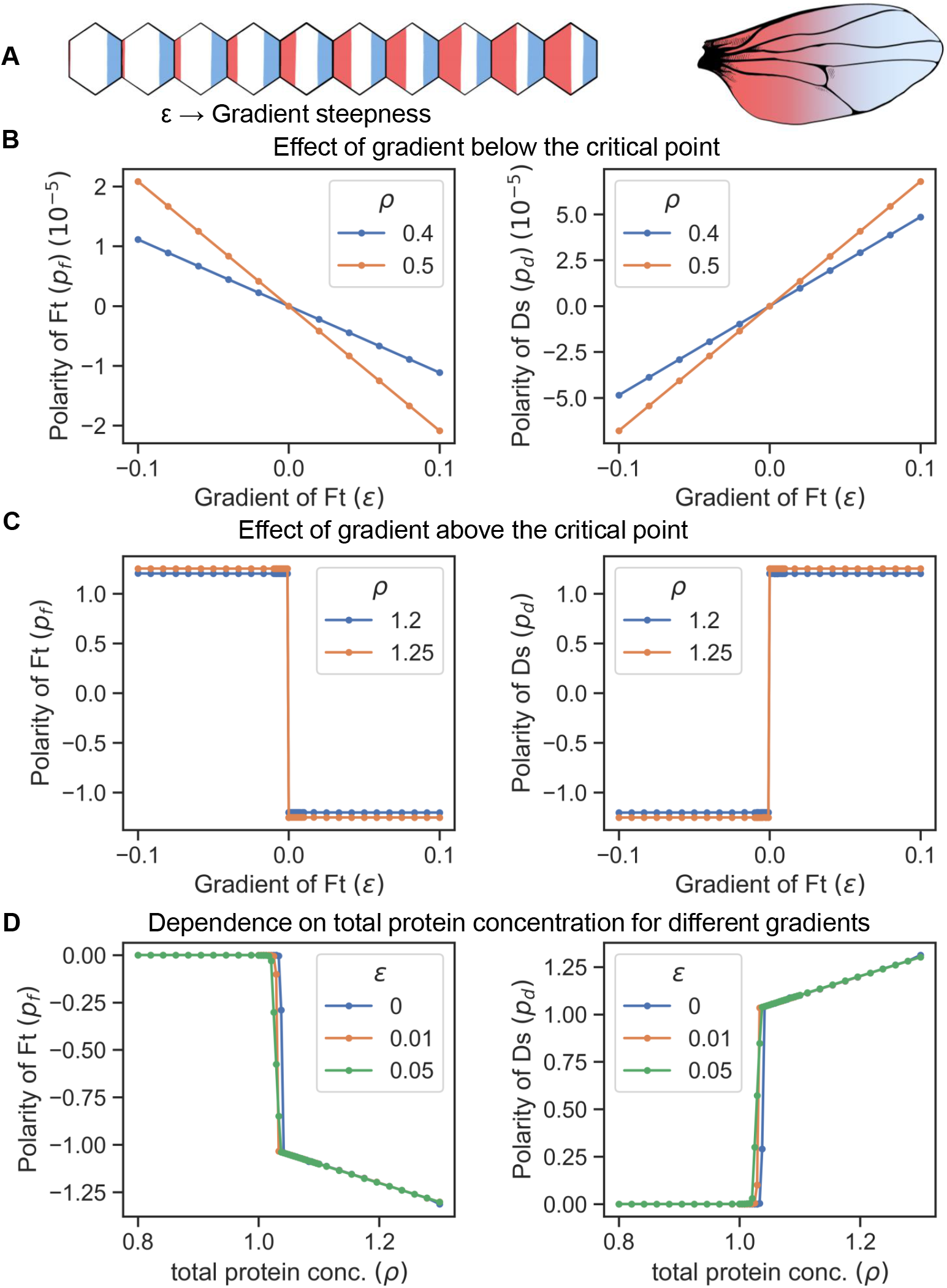
Gradient can polarize the tissue and decide its direction. **A)** Schematic to describe the model of gradient of expression of Ft (red) in a tissue. **B)** Polarity of Ft (*p*_*F t*_) and Ds (*p*_*Ds*_) as a function of gradient of Ft (*ϵ*), when total protein concentration (*ρ*) is below the critical point. The tissue is polarized due to the presence of gradient and the direction is opposite to that of the gradient. **C)** Polarity of Ft (*p*_*F t*_) and Ds (*p*_*Ds*_) as a function of gradient (*ϵ*) when total protein concentration (*ρ*) is above the critical point. Here, the gradient decides the direction of the polarity. **D)** Polarity of Ft (*p*_*F t*_) and Ds (*p*_*Ds*_) as a function of total protein concentration (*ρ*) for different values of gradient (*ϵ*)

We incorporate the linear gradient in Ft expression such that the total protein concentration of Ft varies in the tissue as

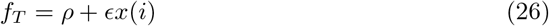

with *ϵ* ≪ *ρ* and *ρ* is the concentration of Ft at some reference location in the tissue. Concentration of Ds is taken to be uniform throughout the tissue and is equal to *ρ*. Here, *ϵ* decides the steepness or the strength of gradient in the tissue and *x*(*i*) is the position of the cell (having index *i*) in the tissue. We studied the effect of expression gradients in both 1D and 2D. In both cases the results are largely similar (**Fig** 5 and **Fig** S5).

If *ϵ* is positive then, for each cell, the neighbour towards right has more total concentration of Ft and the one on the left has lesser concentration as shown in **Fig** 5 A and **Fig** S5 A. We present the results from the one-dimensional model in the main text; our findings from the two-dimensional model are presented in the supplementary information.

Recall that, in the absence of any gradient, the tissue is unpolarized if the total concentrations of Ft and Ds are below their critical values. This, however, is not the case when a gradient of expression is present. In the presence of a gradient, the tissue is polarized even if total protein concentration is below threshold critical (*ρ < ρ*_*c*_) (**Fig** 5 B and **Fig** S5 C). However, the magnitude of polarization is small in this case if the gradient is small. The magnitude of polarity in Ft and Ds depends on the steepness *E* of the expression gradients and also on *ρ*. Interestingly, polarization of Ft (*p*_*f*_) is in the direction opposite direction to that of the gradient. We can understand this observation in the following way. For a given cell, if the neighbour on the right has more Ft concentration than the one on left, then Ds will localise more towards right because of the inter-cellular heterodimer formation. This process will happen in all cells, Ft from the neighbour towards right of each cell will start localising towards left side of the neighbour, to form heterodimer with Ds. This feedback loop amplifies their initial differences over time and will result in eventual localization of Ft towards the left side of all the cells, and Ds on the right side.

If the total protein concentration of Ft and Ds is above the threshold protein concentration, the tissue polarizes due to inter-cellular interactions but the gradient decides the direction of polarization. Due to the combined effect of gradient and spontaneous polarization, in this scenario, the cells get strongly polarized (compare the values on the *y* axes of Figs. 5B and 5C). Polarization of Ft is again opposite to the gradient direction. The physics of spontaneous polarization implies that an arbitrarily weak gradient decides the direction of polarization and the tissue then polarizes due to the feedback loop of inter-cellular interactions (**Fig** 5 C and **Fig** S5 D).

We studied the effect of noise in protein kinetics to understand how noise disrupts the directional cue provided by gradient(**Fig** S4 A). We calculated the polarity of Ft (*p*_*f*_) and Ds (*p*_*d*_) as function of gradient of Ft (*ϵ*) for different noise amplitudes (*η*). We observe that the gradient still decides the direction polarity in presence of noise, but noise also leads to some cells polarizing in the direction opposite to the majority. resulting in a reduction of overall polarity in the tissue.

Next, we study how a gradient affects the robustness of polarization in presence of noise. We calculated the polarity of Ft (*p*_*f*_) and Ds (*p*_*d*_) as function of noise amplitudes (*η*) for different gradient steepness of Ft (*ϵ*)(**Fig** S4 B). We observe that higher the gradient, higher noise amplitude is needed for the system to lose the polarization. Therefore, the gradient helps in maintenance of the polarized state and system with higher gradient is more robust to perturbations. Finally, we study the effect of total protein concentration (*ρ*) on polarization of Ft (*p*_*f*_) and Ds (*p*_*d*_) for various gradient amplitudes (*ϵ*) (**Fig** 5 D and **Fig** S5 E).An increase in the gradient shifts the transition between weak and strong polarization of the cells to smaller values of *ρ*.

To understand analytically how a gradient facilitates polarization, we use the continuum model [38]. We solve the system for the case where total protein concentration is below the critical protein concentration and *ϵ* ≪ *ρ*. For this we get for the steady state

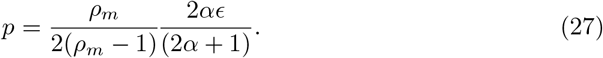

This equation tells us that the tissue will always be polarized in presence of the gradient. The slope of polarization (*p*) vs (*ϵ*) is determined by (*ρ*_*m*_) which in turn is determined by the protein binding rate (*α*) and total protein concentration of Ds (*ρ*). We also performed the stability analysis which tells us that the polarization is stable against non-homogeneous perturbations in the presence of a gradient ([38]).

These results show that expression gradients of the proteins in the PCP also play other roles in addition to providing global cue to align PCP direction with tissue axis. A gradient not only helps in polarity establishment when overall protein levels are low but also stabilizes the polarized state as well.

### Down-regulation of Ft from a region in absence of gradient leads to exponentially decaying isotropic polarity defects

In experimental investigations, in order to identify the role of the protein interactions and their non-cell autonomous effects, the protein expression levels are perturbed from a select regions of the tissue [21]. In one such experimental strategy Ft or Ds are downregulated or upregulated from a region of the tissue and its fall out is observed in the cells of that region as well as outside of it. In order to recapitulate this scenario *in silico*, we set the total concentration of Ft in a region of the 1D tissue (the left half) to be zero as shown in **Fig** 6A while keeping it to be uniform and nonzero in the right half of the tissue. On the other hand the the total protein concentration of Ds is kept uniform and non-zero. To deal with the discontinuity at the deletion boundary we modify the equations by adding an extra term in them as described here:

**Figure 6.**
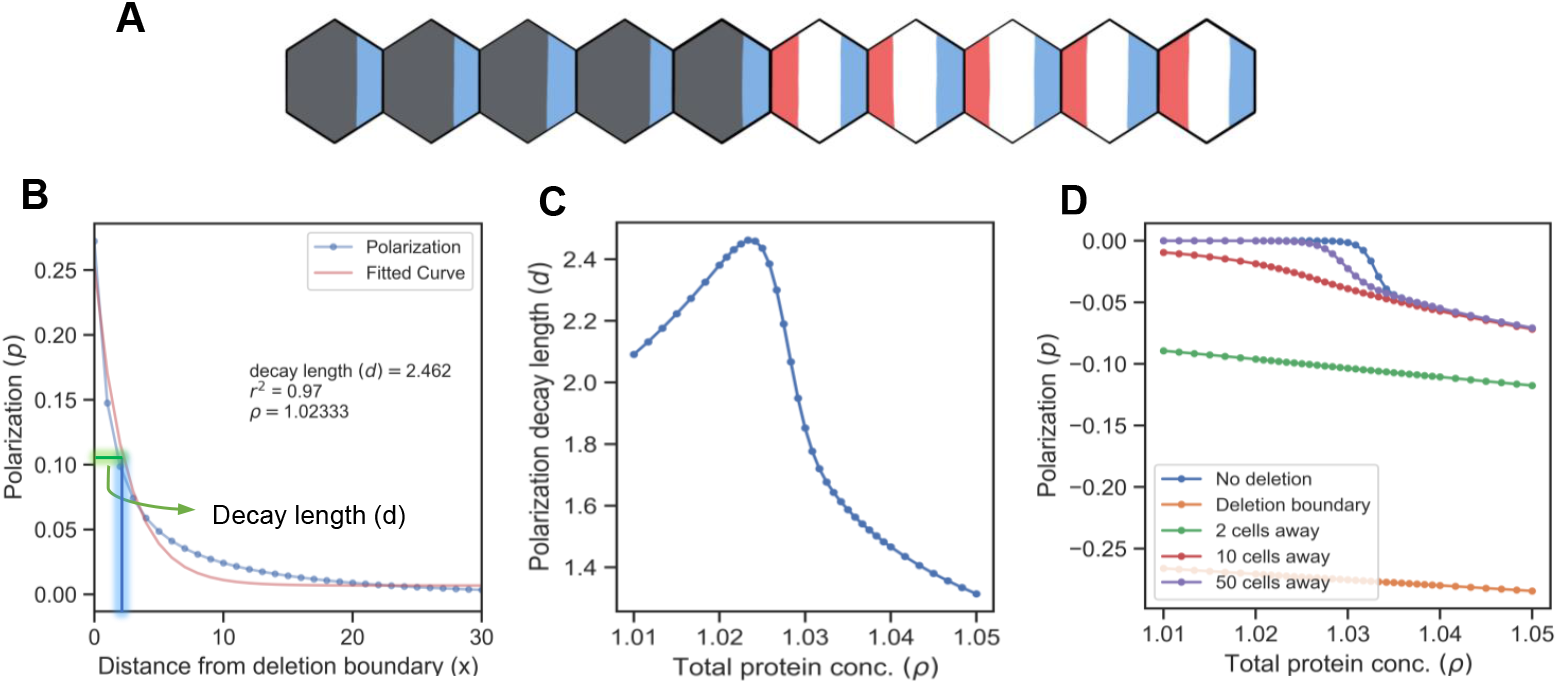
Deletion induces a polarity which decays exponentially. **A)** Schematic to describe the model of deletion of a gene from one side of the tissue. Ft (red) is deleted from the left side of the tissue. **B)** Plot of polarity (*p*) of each cell as function of distance from the deletion boundary. (*x*) is measured from the deletion site and corresponds to the number of cells. Polarization decays exponentially. **C)** Decay length of polarization as a function of total protein concentration. Exponential decay was fit to polarization of 50 cells after the deletion boundary. **D)** Polarization at different points in the tissue (for different distances from the deletion boundary). No deletion is the control case in which there is no deletion of any gene from the tissue.

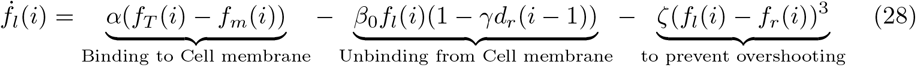

Here, the third term *ζ*(*f*_*l*_(*i*) − *f*_*r*_(*i*))^3^ encodes saturation, making sure that the concentrations do not grow without bound This term is not included except where the unpolarized state is unstable. It can safely be set to zero otherwise without altering the results.

When the total protein concentration is above the critical concentration, we observe that deleting Ft from a region induces a polarization which decays as we move away from the deletion boundary (**Fig** 6 B). Ft from the cell towards right of the deletion boundary binds to the Ds from the cell towards left of the deletion boundary. This leads to more Ft localising towards left and Ds towards right. As we move away from the deletion boundary, the effect of this inter-cellular interactions decays. We fitted the decay of polarity to a decaying exponential function and calculated the decay length of the polarization, i.e., the length or number of cells after which the polarization reduces to *e*^*−*1^ of its value at the deletion boundary. Next, we calculate this decay length as a function of total protein concentration (**Fig** 6 C). Finally, we plot the polarization as a function of total protein concentration at several distances from the deletion boundary. Specifically we calculate the polarization as function of total protein concentration at the deletion boundary and at points which are 5,10,50 cells away from the deletion boundary and for the case when there is no deletion in the tissue (all cells have equal Ft and Ds). As expected, as you move away from the deletion boundary (50 cells away), the tissue behaves like the case when there is no deletion at all. (**Fig** 6 D). To understand why polarity is formed at the deletion boundary and why it follows an exponential decay profile we use our continuum model which can be solved analytically (see Supporting Information for details). We get

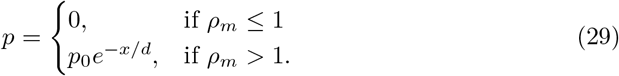

where *d* = (*ρ*_*m*_*/*2)[(2*α* + 1)(*ρ*_*m*_ − 1)]^*−*1*/*2^ is the decay length of polarity and *p*_0_ can be determined by the boundary condition at *x* = 0. The continuum model also predicts the exponential decay in polarity which is in agreement with our discrete model results.

Next, we deleted or up-regulated Ft from a circular region of the tissue in the 2D model. The total protein concentration of Ft is same as that of Ds throughout the tissue but the circular region where it is either zero or a fixed high value (**Fig** 7 A). Our aim is to account for the experimental observation that cells near the deletion boundary are polarized towards or away from the deletion region if a gene is deleted or up-regulated in that part respectively (**Fig** 7 B).

**Figure 7.**
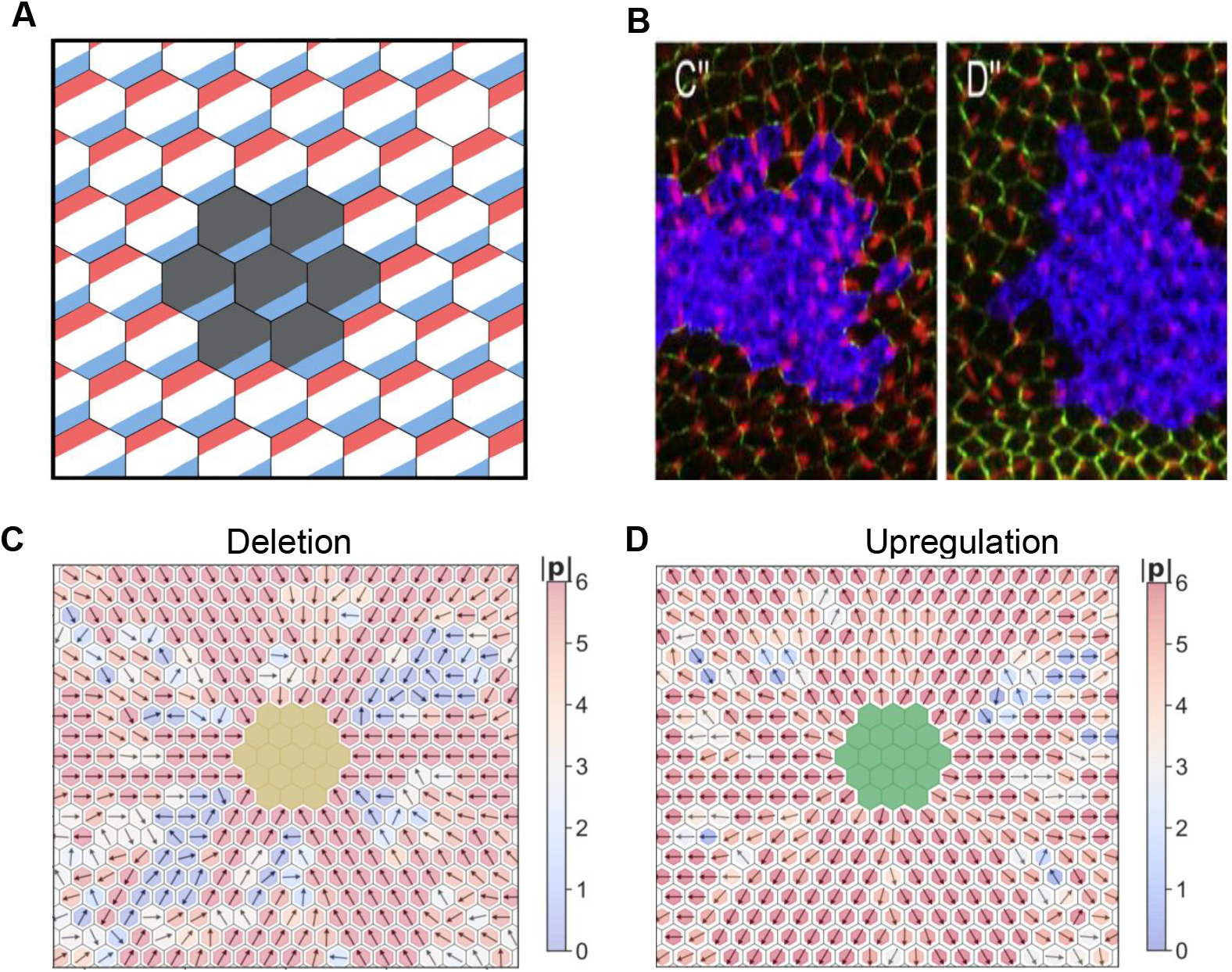
Deletion and upregulation in a 2D layer of tissue. **A)** Schematic to show deletion of Ft (red) from cells lying in the circular region (grey) while other cells have equal concentration of Ds and Ft. **B)**Experimental images for deletion/upregulation taken from Chen et al.(2008) [44]. Cells around the deletion regions (blue) are polarized in the direction. **C)** Polarization (**p**) when Ft is removed from a portion of the tissue (yellow) and total protein concentration (*ρ*) is above the critical point. **D)** Polarization (**p**) when Ft is removed from a portion of the tissue (yellow) and total protein concentration (*ρ*) is above the critical point. **E)** Polarization when Ft is up-regulated in a portion of the tissue (green) and total protein concentration (*ρ*) is below the critical point. **F)**Polarization when Ft is up-regulated in a portion of the tissue (green) and total protein concentration (*ρ*) is above the critical point.

When Ft is deleted from a region, and total protein concentration of Ft and Ds is below the threshold value, the cells in the vicinity of the deletion region are polarized radially towards the deletion region. But as the effect of the deletion decays away, the magnitude of polarization in cells also goes down (**Fig** 7 C, **S Video** 3).

If the total protein concentration is above the threshold value, the most of the cells in the tissue, polarize radially in the direction of the deletion region (**Fig** 7 D). The magnitude of polarization is also greater in this case compared to the case where total protein concentration is below the critical point.

When Ft is up-regulated in the region, the results are opposite of the deletion case. Now cells are polarized away from the upregulation region (**Fig** 7 E and F). We can understand these results by simply thinking of the feedback loop. If Ft is deleted from a region, the Ft from the surrounding region will make a heterodimer with the Ds of the cells inside the deletion boundary. This will in turn result in localisation of Ds from the surrounding region moving away from deletion region which result in Ft from the next circular layer of cells to localise inwards. This feedback loop will result in polarization of the tissue in the direction of deletion. Similar process happens in the case when Ft is up-regulated except that the directions will be reversed.

### Presence of gradient and local down-regulation explains the domineering non-autonomy

Finally, we tried understanding one of the most interesting features of PCP. We incorporate both gradient in concentration of Ft and deleted it from a central region in the tissue (**Fig** 8 A). We aim to understand the phenomenon of domineering non-autonomy [33, 45] where disruption in protein levels of a mutant cell in the tissue affects the polarity of neighbouring wild-type cells (**Fig** 8 B). When we simulated the system, we observe that the cells decide the direction of polarity depending on relative proximity to the deletion region, and strength and direction of gradient. For the cells close to the deletion region, effect of deletion is strong, and they polarize according to the deletion. On one side of the deletion, the gradient supports this polarization and on the other side it opposes the polarization. On the side where it opposes the polarization, cells polarize in direction opposite to the deletion region if they are far enough from the deletion region (**Fig** 8 C-D,, **S Video** 4).

**Figure 8.**
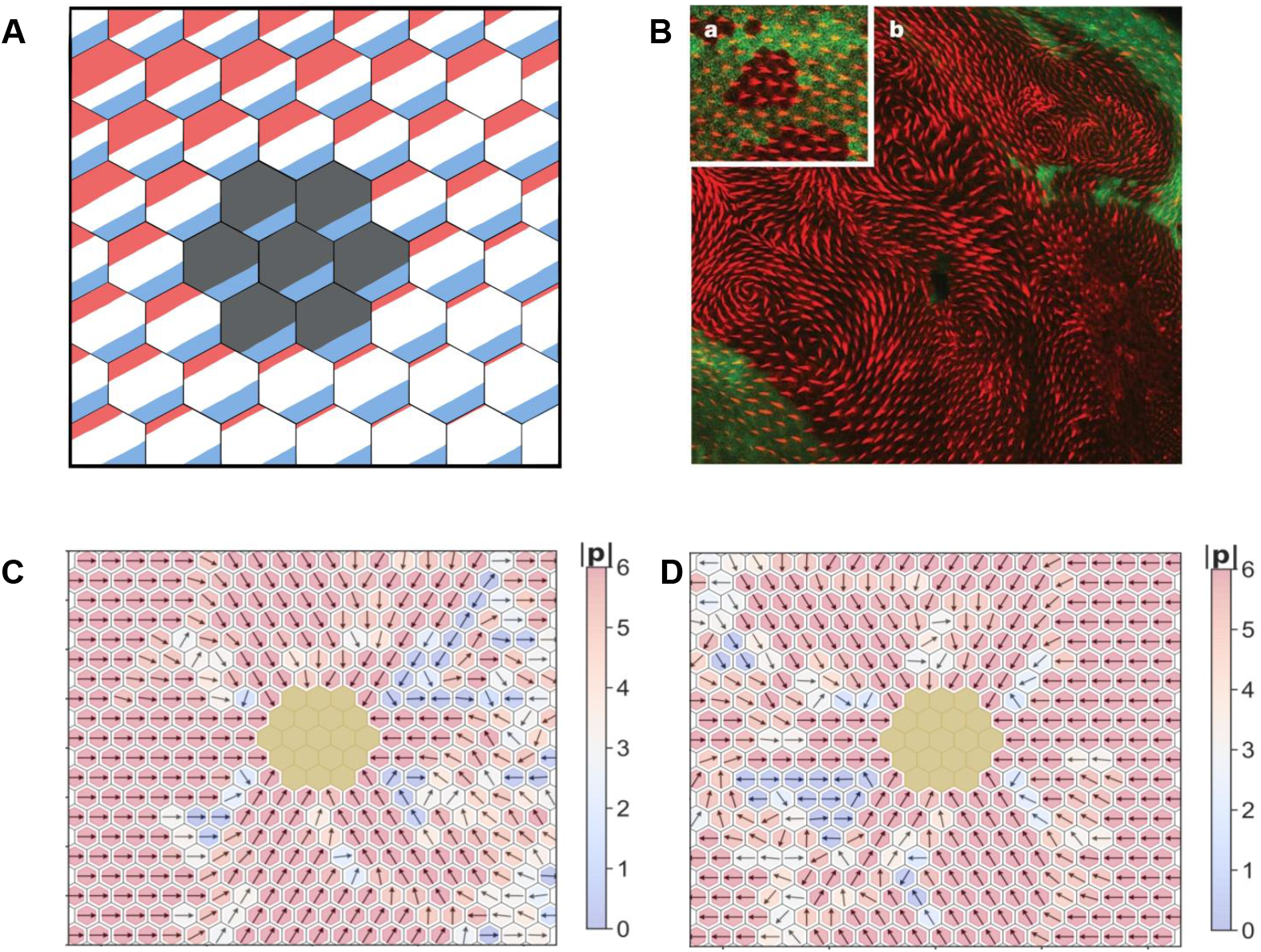
Deletion and gradient can explain domineering non-autonomy. **A)** Schematic to explain the model of tissue where Ft (red) has been deleted from cells that lie in the circular region (red) and there is a gradient in its expression. **B)** Experimental image taken from [21] to demonstrate domineering non-autonomy **C)** Polarization (**p**) in the tissue when gradient (*ϵ*) is in the positive direction. The polarity is disrupted only on left side of the deletion region.**D)** Polarization (**p**) in the tissue when gradient (*ϵ*) is in the negative direction. The polarity is disrupted only on right side of the deletion region.

## Discussion

The minimal discrete model presented here offers a useful framework for combining the role of global (expression gradients) and local (heterodimer formation) interactions in PCP and studying their collective effects. Our framework builds upon the discrete modeling of global/local modules [12, 25] and other phenomenological models [27, 31]. Our model stands apart from the previous models like [27] because here we have modelled the protein interactions instead of heterodimer interactions. Since experimental studies haven’t confirmed the interaction of heterodimers per se, the current model based on protein-protein interactions (as opposed to heterodimer-heterodimer interactions [27, 32] which can be considered an outcome of protein-protein interactions) provides a better motivated framework to understand PCP. Due to its minimalist nature (with only three parameters) the model presented here also differs from other protein-protein interactions based studies, such as [12, 25, 28], which involve high number of model parameters. In this study, we have also studied the effect of noise in total protein concentration (quenched noise) in the cells. These spatial heterogeneities in the protein expression levels lead to deterministic dynamics resulting in static polarization patterns as opposed to dynamic patterns due to the time dependent noise in the kinetics of protein-protein interactions (akin to the one studied in [30]).

Some earlier studies like [30, 31] have invoked the similarities between XY model of ferromagnetism and PCP formation. This has helped in understanding the formation of swirling patterns and the effect of gradient in the tissue level polarization.

In this model, we have taken account of simple mechanistic details which are common for both core and global modules. Thus it helps in understanding the properties of both the modules and the predictions from our model can be tested experimentally. For instance, the exponential decay in polarization near a clone boundary, as predicted by the model, can be measured experimentally. Another experimentally testable prediction from the model is the effect of the protein expression gradients on the domineering non-autonomy where loss of the two proteins, in the presence of expression gradient, can affect the two sides of the clone differently. On the other hand, in absence of a gradient, we observe a disruption in polarity in areas adjacent to the mutant clone but the disruption is isotropic in the sense that the polarization direction near clone boundary is perpendicular to it. This prediction can also be tested experimentally by perturbing gradients and generating mosaic patterns of deletion or overexpression of specific proteins.

Our model also offers advantages over other mechanistic discrete models which have several free parameters making it difficult to analyze [46]. After non-dimensionalizing, we are left with just three free parameters: protein binding rate (*α*), total protein concentration of Ft and Ds (*ρ*) and gradient steepness (*ϵ*) which are very easy to control and study.

In fact, we could also analyze the discrete model in the continuum limit which gives some valuable insights which are validated systematically over a wide range of parameter values in discrete version. Conceptually, the first two parameters (*α* and *ρ*) correspond to the “strength” of relay of PCP “signal” from one cell to another; while the last one (*ϵ*) modulates the direction of signal propagation and offers robustness against any stochastic perturbations. In absence of gradient, the polarity is still established in individual cells but is not necessarily coordinated at the tissue level. When the local interaction is weak (total protein concentration is low), the global interaction is responsible for formation and direction of polarity. On the other hand, when local interaction is strong, gradient helps in providing the directional cue and maintenance of polarity in presence of perturbation. Our prediction of gradient providing the robustness against perturbation can be experimentally verified by constructing clones with different gradient strength and measuring polarization in different cases. The clones where strength of gradient is less, we should see more cells which are misaligned compared to the case where the gradient strength is more. This prediction can be extended to claim that polarity establishment in individual cells is not disrupted, but the synchronization is lost. One special case of such disruption is formation of swirling patterns seen in 2D; which the model is able to recapitulate. Thus, the minimal model can explain diverse experimental observations seen in many biological systems; suggesting common design principles at play.

In this model we have only considered the most simplistic and common interactions seen in the proteins involved in PCP. To keep the model simple and widely applicable, we have not taken into account various inter-cellular and intra-cellular interactions which are known in the literature. In our model, we have considered the small additive noise in protein kinetics and total protein concentration. The underlying the assumption here is that the average copy number of proteins are much larger than the levels of noise. But for the tissues where this assumption is not valid, the model needs to be modified [32].

One of the major limitation of current model is that it doesn’t account the change in lattice structure due to cell death, birth and growth and epithelial flows [47]. Modelling studies like [32] discusses the effect of cell shape changes on polarity formation. Future work needs to incorporate these factors in our discrete modelling framework.

## Data Availability Statement

All codes used in this study are publicly available on the GitHub page of D.S. (https://github.com/Divyoj-Singh/Planar_cell_polarity).

## Acknowledgments

We would like to acknowledge Atchuta Srinivas Duddu for the schematics. D.S. is supported by KVPY fellowship awarded by Department of Science and Technology (DST), Government of India. MKJ is supported by Ramanujan Fellowship (SB/S2/RJN 049/2018) awarded by the Science and Engineering Research Board (SERB), DST, Government of India. SR acknowledges support from a J C Bose Fellowship of the Science and Engineering Research Board, India. MSR would like to acknowledge the financial support provided by the seed grant from IIT Hyderabad.

## Supporting Information

### Continuum limit of the one-dimensional model

This section has been adapted from [38]. The reader is encouraged to refer to the original work for more details and the continuum counterpart of two-dimensional model.

In 1D we can describe the protein distributions by *f*_*l*_(*x*) (amount of *Ft* on left edge of cell), *f*_*r*_(*x*), *d*_*l*_(*x*) and *d*_*r*_(*x*). Here *x* is the cell coordinate. The equations are (*x* (position) and *t* (time) dependencies are not written explicitly)

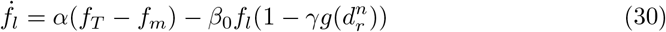

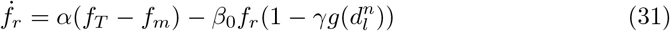

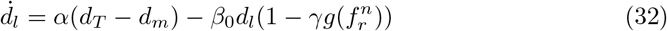

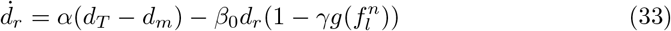

where *f*_*l*_(*x*), *f*_*r*_(*x*), *d*_*l*_(*x*), *d*_*r*_(*x*) represent the amount of Ft and Ds localized on the left or right edge of the cell, 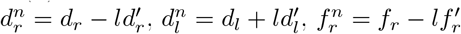 and 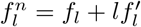 represent the concentration of Ds and Ft in the neighboring cell edges. For simplicity we have taken *g*(*c*) = *c* which is the leading order behavior of Hill’s function 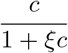 which is usually used for the modeling of protein-protein interactions. ()^*′*^ and 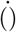 denote the spatial and time derivatives respectively, and *f*_*m*_(*x*) = *f*_*l*_(*x*) + *f*_*r*_(*x*) (total membrane bound Ft in a cell at location *x*) and *d*_*m*_ = *d*_*l*_ + *d*_*r*_. The terms *f*_*T*_ and *d*_*T*_ respectively stand for the amount of total (cytoplasmic + membrane bound) Ft and Ds proteins in the cell and can be dependent on *x*, the cell location in the tissue.

We can define *p*_*f*_ = *f*_*r*_ − *f*_*l*_ and *p*_*d*_ = *d*_*r*_ − *d*_*l*_ to be the asymmetry (or polarity) in the localization of Ft and Ds on cells edges. In order to ensure that the protein levels do not become negative at any of the cell edges, this definition of protein asymmetry requires |*p*_*f*_ | ≤ *f*_*m*_ and |*p*_*d*_| ≤ *d*_*m*_. Using the above equations, we can write the equations for *f*_*m*_, *d*_*m*_, *p*_*f*_ and *p*_*d*_ in the non-dimensionalized form as

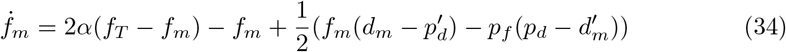

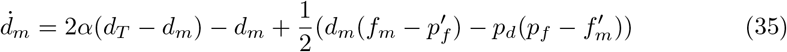

and

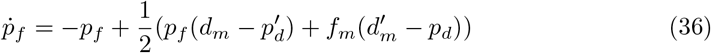

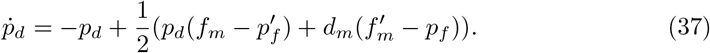

Now, we will deal with the different cases in detail.

### Uniform protein expressions

For the uniform expressions of proteins *Ft* and *Ds* in the tissue, we have *f*_*T*_ (*x*) = *d*_*T*_ (*x*) = *ρ* (a known constant value). If we assume the total membrane bound Ft and Ds to be equal to each other, that is *f*_*m*_ = *d*_*m*_ = *ρ*_*m*_ (an unknown), and we obtain

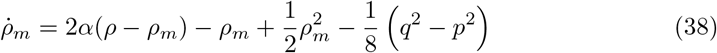

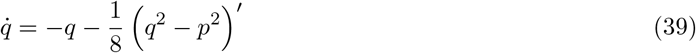

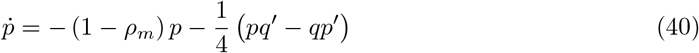

where *p* = *p*_*f*_ − *p*_*d*_ and *q* = *p*_*f*_ + *p*_*d*_. A comparison of these equations for 1D and 2D (presented in [38]) with those for the active matter shows that this system is a generalization of the field theory for the active matter [48, 49]. This outcome is in contrast to the previous works where the PCP dynamics has been considered to be equivalent to the ferromagnetism [30, 31].

These equations show that for very small values of *ρ*_*m*_ the only homogeneous solution (where all spatial derivatives vanish) is *q* = 0 and *p* = 0 at the steady state (unpolarized state). For *ρ*_*m*_ *>* 1, however, we get *q* = 0 and *p >* 0 (polarized state) which imply polarization of two proteins in opposite directions. For very large values of *ρ*_*m*_ the system can also have *q >* 0 which implies unequal polarization of the two proteins in the cells. In this work we do not consider such high levels of *ρ*_*m*_.

### Non-uniform protein expressions

In order to study the effect of gradient, we consider a weak linear [50] gradient of the expression of Ft such that *f*_*T*_ = *ρ* + *ϵx* (with *ϵ* ≪ *ρ*). We assume the following ansatz 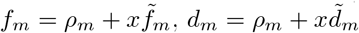 with homogeneous polarization levels in the tissue. For this case, we get

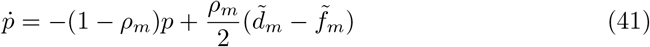

where

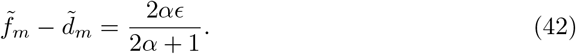

In equation (41) above we have written only the leading order terms on the right hand side. For the steady state, we obtain for *ρ*_*m*_ ≪ 1

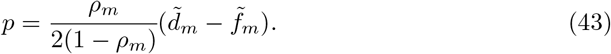

This shows that in case of a tissue-level gradient expression of one of the two proteins, the tissue is always polarized, and that too in a particular direction which depends on the direction of the gradient.

### Loss of one protein from a region of the tissue

In order to model the loss of Ft from a region of tissue we consider Ft to be absent in the cells at *x <* 0 and focus on the polarization in the cells at *x* ≥ 0. This means we are effectively considering a semi-infinite 1D region of *x* ≥ 0. The loss of Ft in the region *x <* 0 will set the boundary conditions for the unknown variables *f*_*m*_, *d*_*m*_, *p*_*f*_ and *p*_*d*_ at *x* = 0. In this case, we expect *f*_*m*_, *d*_*m*_, *p*_*f*_ and *p*_*d*_ to be *x*-dependent and their derivatives with respect to *x* cannot be ignored. Still, we should get the wildtype behavior as seen above for *x* ≫ 1. We presume solutions of type 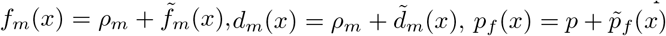 and 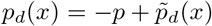 with |*p*| ≪ *ρ*_*m*_.

Substituting these values in the equations (34) and (35) and looking for the steady state solutions gives us

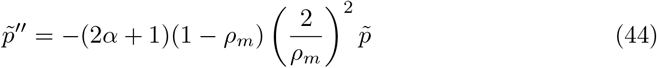

where 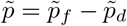 is the excess cell polarization due to the loss of Ft in the left half of 1D tissue. Here we have considered only the leading order terms on the right hand side. Depending on the value of *ρ*_*m*_ this equation can have either exponentially decaying solution (for *ρ*_*m*_ *>* 1) or spatially oscillatory solutions (for *ρ*_*m*_ *<* 1). In order to have a ‘wildtype’ behavior far from *x* = 0 boundary, that is 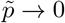 as *x* → ∞, the spatially oscillatory solution becomes unfeasible. This gives us the excess polarization to be

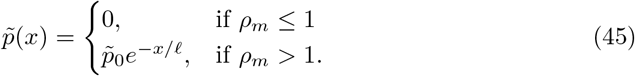

where *𝓁*, = (*ρ*_*m*_*/*2)[(2*α* + 1)(*ρ*_*m*_ − 1)]^*−*1*/*2^, and *p*_0_ can be determined by the boundary condition at *x* = 0. This demonstrates an exponential decay in the magnitude of cell polarization from the boundary of the region where one of the two proteins is not expressed.

## Supplementary Figures

**Figure S1.**
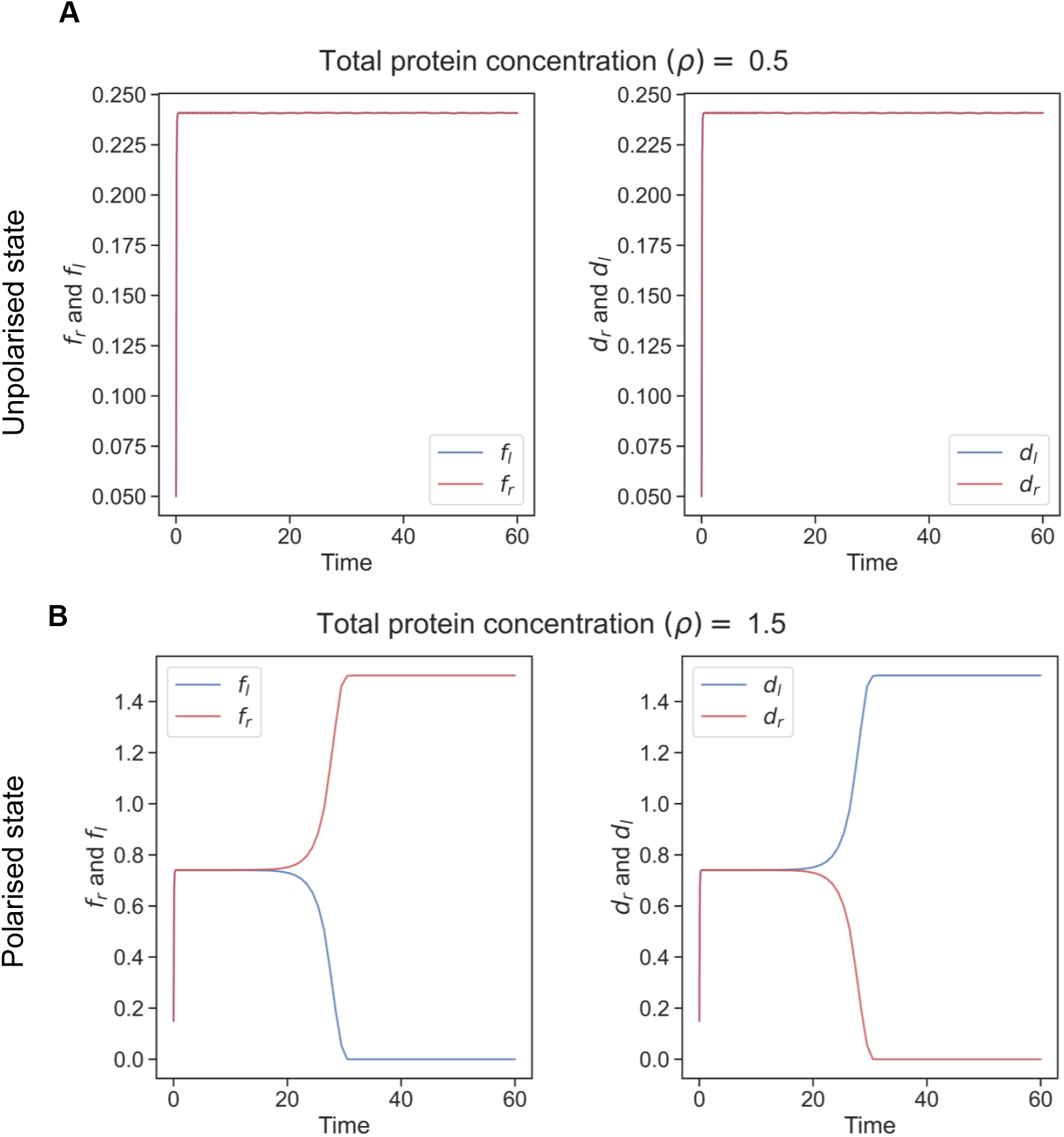
Dynamics in 1D: Level of Ft and Ds in left and right side (*f*_*l*_,*f*_*r*_, *d*_*l*_ and *d*_*r*_) of the tissue as a function of time. **A)** When the total protein concentration of Ft and Ds (*ρ*) is below the critical value both proteins localise equally on both sides of the cell. **B)** But when the total protein concentration of Ft and Ds (*ρ*) is above the critical value, we observe that the proteins localise completely to one side of the tissue.

**Figure S2.**
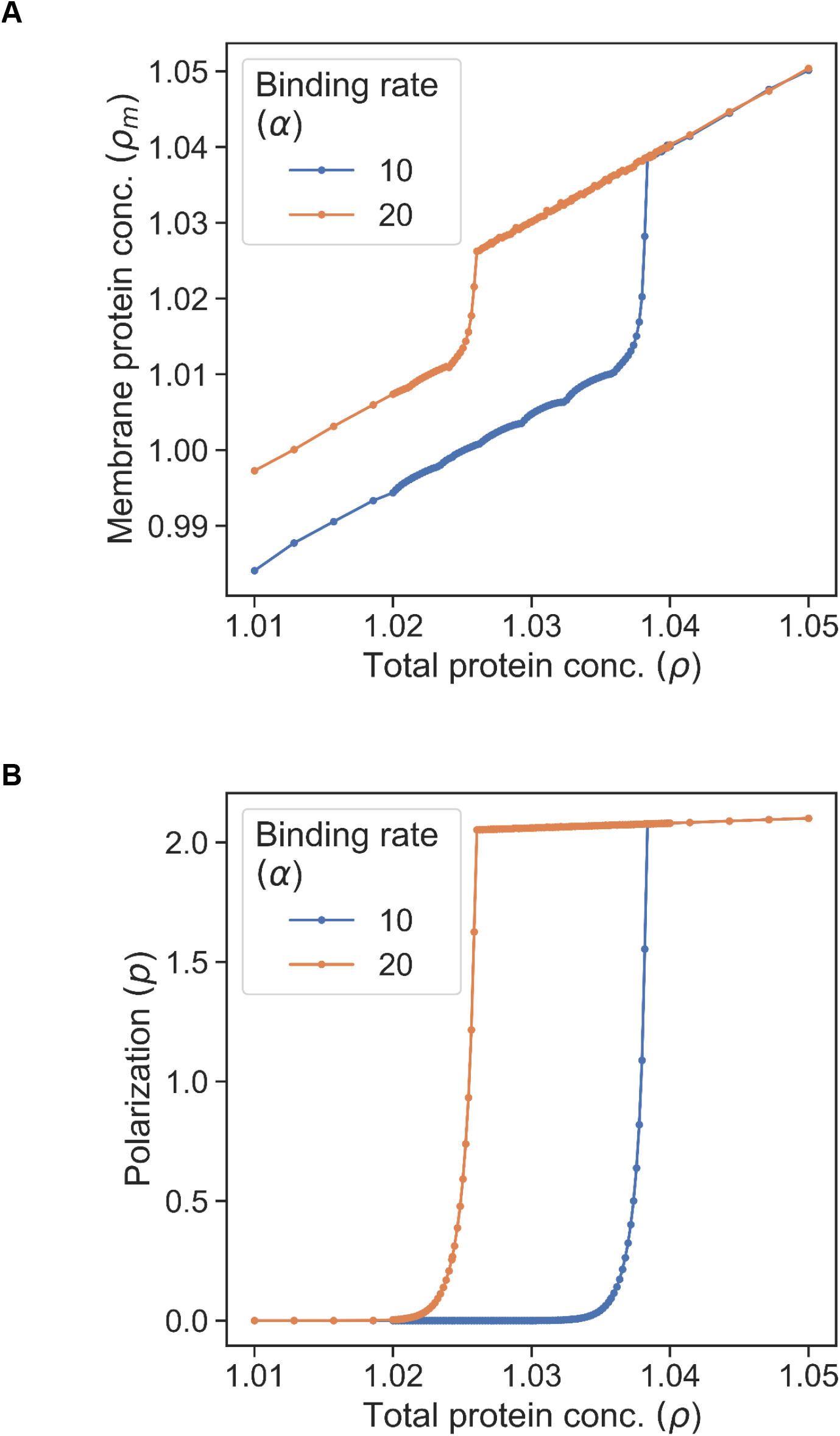
Zoomed plots for Figure 2. **A)** Zoomed plot for membrane protein concentration (*ρ*_*m*_) vs total protein concentration (*ρ*) **B)** Zoomed plot for polarization (*p*) vs total protein concentration (*ρ*). These plots show that the transition from unpolarised state to polarised state is a continuous one.

**Figure S3.**
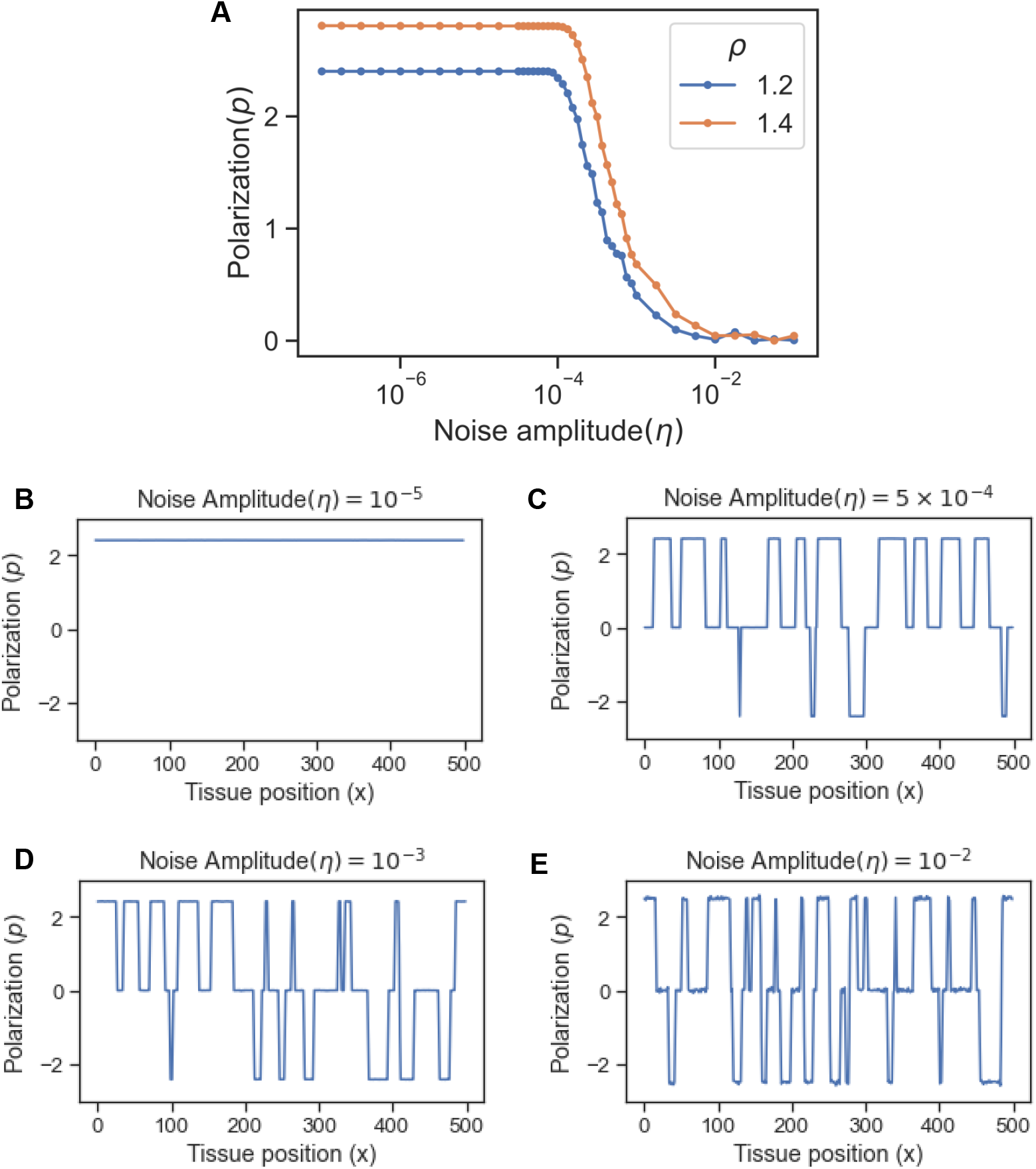
Effect of noise in protein kinetics on polarization. **A)** Polarization (*ρ*) as function of noise amplitude (*η*) for different total protein concentrations (*ρ*). **B)** Polarization (*p*) in the tissue when the level of noise is negligible (10^*−*5^). **C)** Polarization at high amplitude of noise (*η* = 5 ∗ 10^*−*4^), some parts of the tissue are aligned in the opposite direction **D)** At higher noise amplitude (*η* = 10^*−*3^), the average polarity of the tissue reduces **E)** At very high noise amplitude (*η* = 10^*−*^2), the tissue loses overall polarization as different parts of the tissue are aligned in opposite direction.

**Figure S4.**
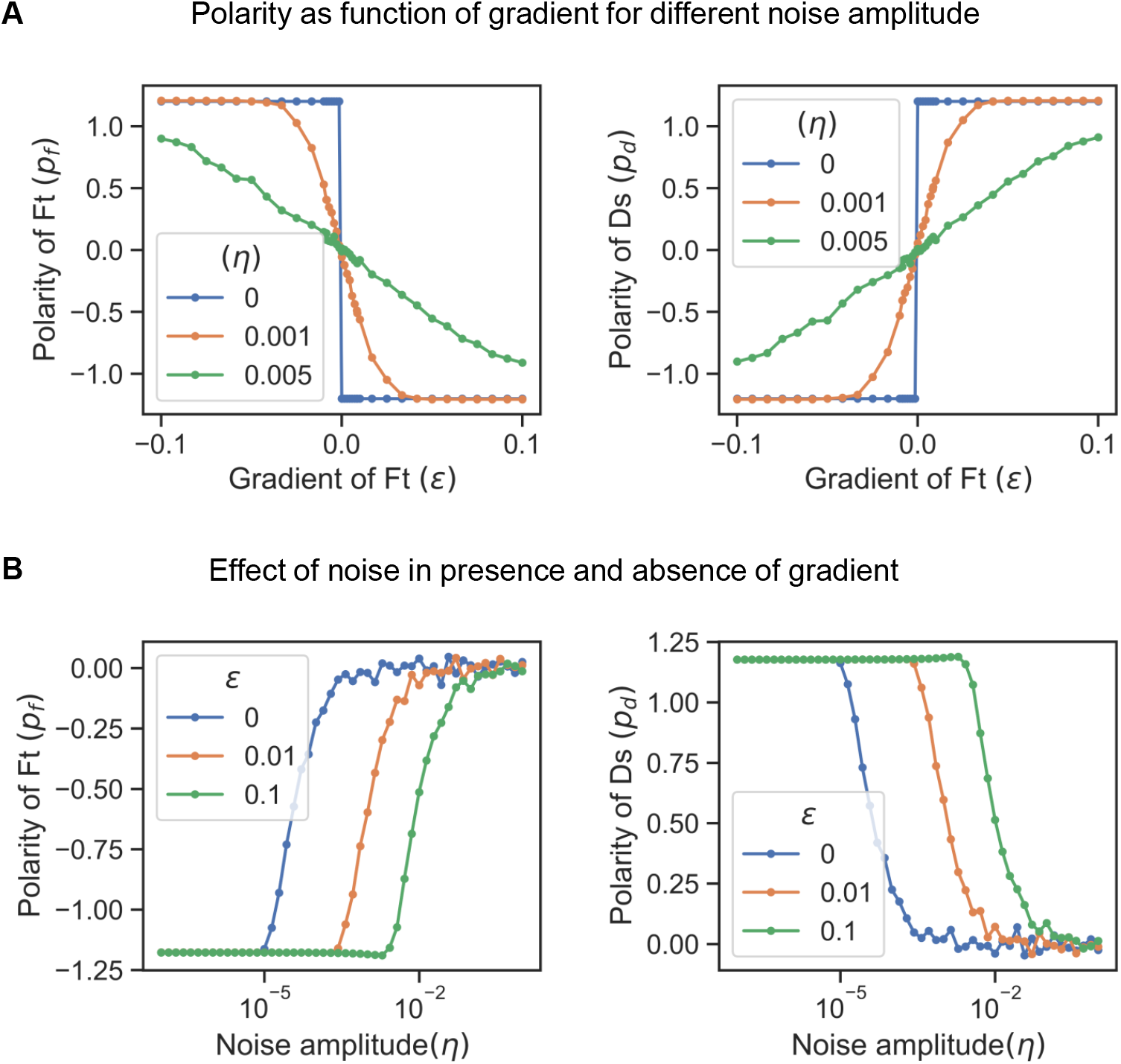
Effect of noise in protein kinetics on the polarization in presence of gradient cue. **A)** Polarization of Ft (*pft*) and Ds (*pds*) as a function of gradient (*ϵ*) in presence of noise (*η* = 0.001, 0.005) and without it (*η* = 0). **B)** Polarization of Ft (*p*_*ft*_) and Ds (*pds*) as a function of noise amplitude (*η*) in presence of gradient (*ϵ* = 0.01, 0.1) and without it (*ϵ* = 0).

**Figure S5.**
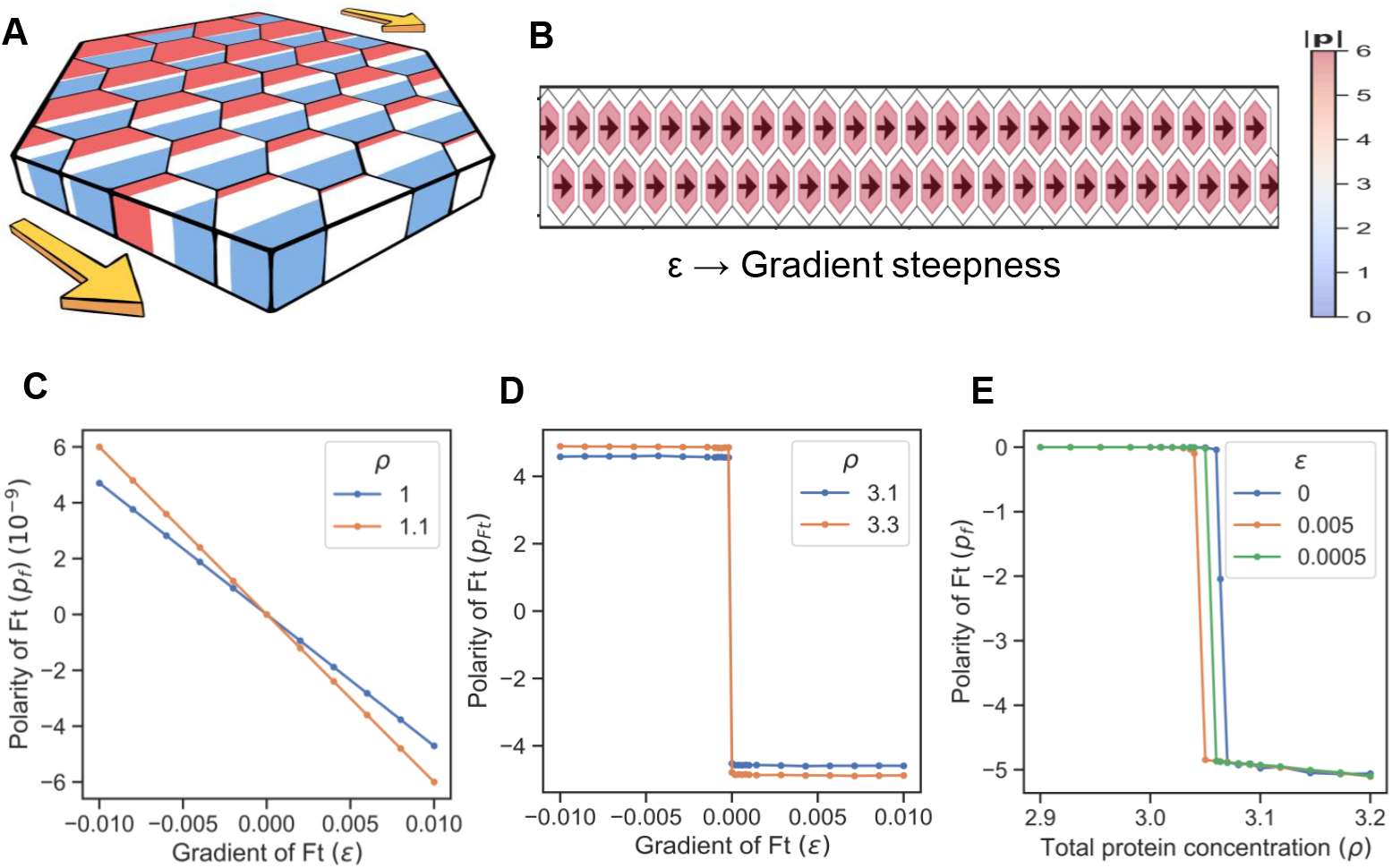
Gradient decides the direction and maintains polarization in 2D: **A)** Schematic of model of a tissue where Ft (red) is expressed in a gradient and Ds (blue) is expressed uniformly throughout the tissue. **B)** Polarization (**p**) in the tissue when gradient is present. **C)** Polarization of Ft (**p**_**f**_) as a function of gradient of Ft (*ϵ*) when the total protein concentration of Ft and Ds (*ρ*) is below the critical point. **D)** Polarization of Ft (**p**_**f**_) as a function of gradient of Ft (*ϵ*) when the total protein concentration of Ft and Ds (*ρ*) is above the critical point. **E)** Polarization of Ft (**p**_**f**_) as a function of total protein concentration of Ft and Ds (*ρ*) for different value of gradient of Ft (*ϵ*).

## Supplementary Videos

1. **Non-homogeneous polarization due to spatial noise in total protein concentration**.: Non-uniform polarization in 2D without gradient. Swirling patterns and vortices are formed. Periodic boundary conditions and initial conditions are such that there is a small misalignment. Total protein concentration in different cells is from a Normal distribution with a mean of 3.2 and a standard deviation of 0.1.
2. **Effect of noise in protein kinetics in a 2 D PCP system:** Additive white noise with mean 0 and standard deviation 0 has been added to the protein kinetics equations. We observe that the tissue gets polarized in a non-homogenous manner and there is fluctuation in polarization of some cells.
3. **Polarization in case of deletions:** One of the genes is deleted from the central part of the tissue. The system gets polarised in the direction of the deletion region.
4. **2-D Polarization in case of deletion with gradient:** One of the proteins is deleted from the central region of the tissue. There is also a gradient across the tissue

